# A half-center oscillator encodes sleep pressure

**DOI:** 10.1101/2024.02.23.581780

**Authors:** Peter S. Hasenhuetl, Raffaele Sarnataro, Eleftheria Vrontou, H. Olof Rorsman, Clifford B. Talbot, Ruth Brain, Gero Miesenböck

## Abstract

Oscillatory neural dynamics are an inseparable part of mammalian sleep. Characteristic rhythms are associated with different sleep stages and variable levels of sleep pressure, but it remains unclear whether these oscillations are passive mirrors or active generators of sleep. Here we report that sleep-control neurons innervating the dorsal fan-shaped body of Drosophila (dFBNs) produce slow-wave activity (SWA) in the delta frequency band (0.2–2 Hz) that is causally linked to sleep. The dFBN ensemble contains rhythmic cells whose membrane voltages oscillate in anti-phase between hyperpolarized DOWN and depolarized UP states releasing bursts of action potentials. The oscillations rely on direct interhemispheric competition of two inhibitory half-centers connected by glutamatergic synapses. Interference with glutamate release from dFBNs disrupts SWA and baseline as well as rebound sleep, while the optogenetic replay of SWA (with the help of intersectional, dFBN-restricted drivers) induces sleep. dFBNs generate SWA throughout the sleep–wake cycle— despite a mutually antagonistic ‘flip-flop’ arrangement with arousing dopaminergic neurons—but adjust its power to sleep need via an interplay of sleep history-dependent increases in excitability and homeostatic depression of their efferent synapses, as we demonstrate transcriptionally, structurally, functionally, and with a simple computational model. The oscillatory format permits a durable encoding of sleep pressure over long time scales but requires downstream mechanisms that convert the amplitude-modulated periodic signal into binary sleep–wake states.

## Introduction

The dynamics of neurons responsible for the homeostatic regulation of sleep, like those of all feedback controllers, are coupled to the dynamics of the controlled object: the outputs of one system serve as inputs to the other. Studies in *Drosophila* have begun to paint a detailed picture of how sleep-inducing neurons sense changes in the physiology of the sleeping and waking organism. Sleep-control neurons^1,2^ with projections to the dorsal layers of the fan-shaped body (dFBNs) estimate sleep pressure by monitoring the flow of electrons through their own mitochondria^3,4^. Sleep loss creates an imbalance between electron supply and ATP demand^3^ that diverts high-energy electrons from the respiratory chain into uncontrolled side reactions with molecular O_2_, producing reactive oxygen species^4^ which fragment the polyunsaturated fatty acyl (PUFA) chains of membrane lipids into short- or medium-chain carbonyls^5^. dFBNs count the release of PUFA-derived carbonyls (and transduce this signal into sleep) in a process that involves an allosteric dialogue between the voltage-gated potassium channel Shaker—a critical determinant of dFBN activity^6^—and its redox-sensitive β-subunit Hyperkinetic^4,5^.

These insights were gained in experiments which clamped certain variables in order to isolate others. For example, to determine how sleep history^2^ or mitochondrial dynamics^3^ alter the current–spike frequency function of dFBNs, the neurons were driven with fixed current patterns; to resolve how cellular redox chemistry^4^, lipid peroxidation products^5^, or arousing dopamine^6^ modulate particular ion channels, the membrane potential of dFBNs was stepped between fixed voltages. How the unclamped dFBN population responds to changes in sleep pressure, and how these responses alter the sleep–wake state of the organism, therefore remains unknown.

## Results

### dFBN slow oscillations

To visualize unperturbed dFBN ensemble activity *in vivo*, we expressed the genetically encoded Ca^2+^-sensor GCaMP6f (GCaMP in brief) under the control of *R23E10-GAL4* and imaged the dendritic tufts of dFBNs by two-photon microscopy. Dendritic fluorescence displayed prominent Ca^2+^ transients, which repeated rhythmically at 0.2–1 Hz (Fig. 1a, Supplementary Video 1). In analogy to the nomenclature in mammals^7^, we refer to activity in the frequency band of 0.2–2 Hz as delta or slow-wave activity (SWA).

**Figure 1.**
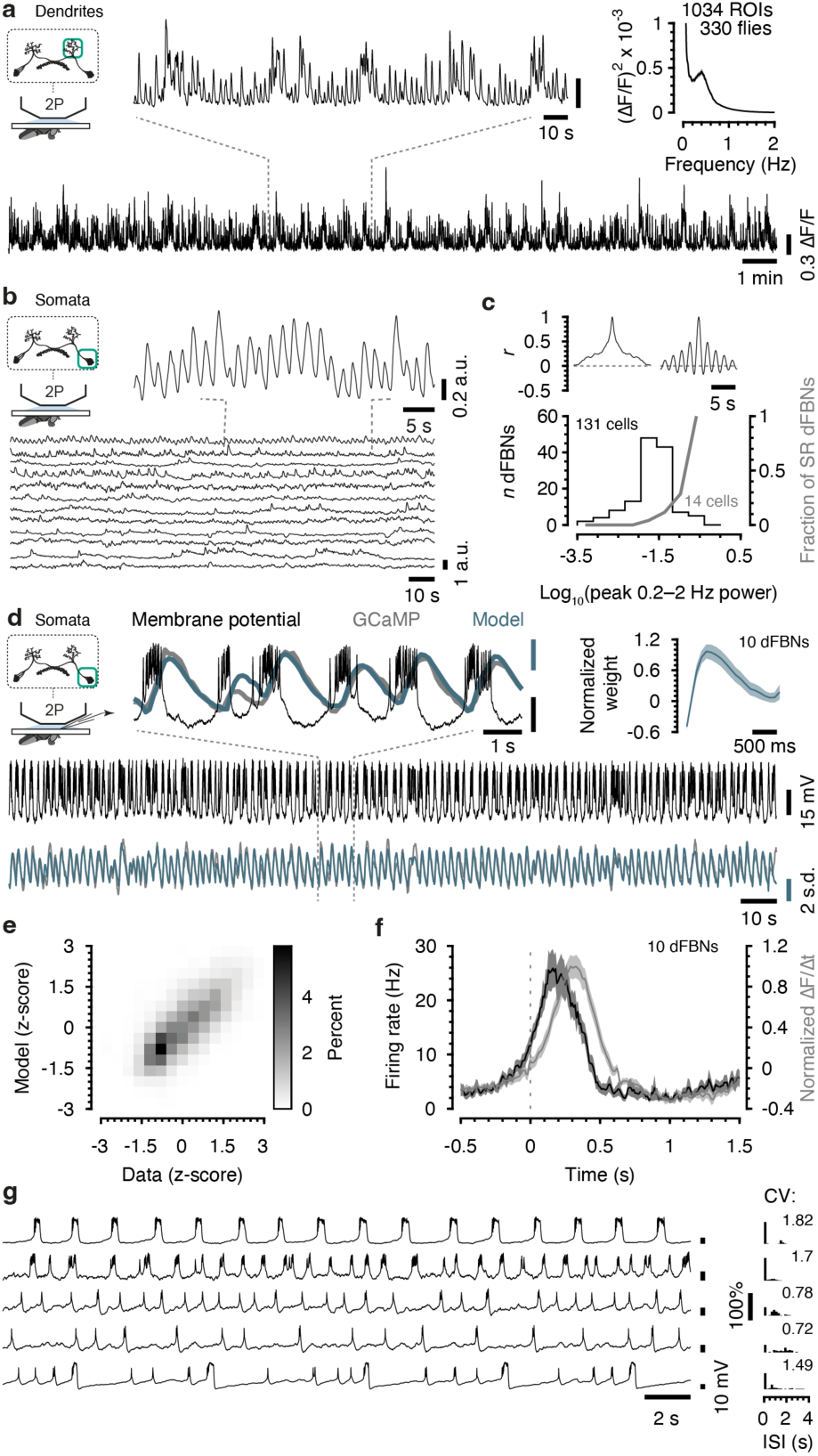
A subset of dFBNs generate slow-wave activity. **a**, Imaging of dFBN dendrites. The example shows 180 s of a GCaMP trace spanning ∼22 minutes (bottom) at expanded x- and y-scales on top. Top right, power spectrum. **b**, Simultaneous imaging of 12 dFBN somata in one hemisphere (intensity-normalized fluorescence). A portion of the uppermost trace is shown at expanded x- and y-scales on top. **c**, Top, GCaMP fluorescence auto-correlograms of the uppermost (right) and lowermost (left) dFBNs in **b**. Bottom, log-scaled distributions of 0.2–2 Hz power among 131 dFBNs in 17 hemispheres of 13 flies (black) and fraction of dFBNs with a slow rhythm (SR dFBNs) in each bin (grey). Prominent periodicity in the auto-correlogram with a wide amplitude swing during the first period was diagnostic of SR dFBNs. **d**, Simultaneous imaging and patch-clamp recording from dFBN somata. Black, grey, and blue traces represent recorded voltages and z-score-normalized measured and predicted GCaMP traces, respectively, shown at expanded x- and y-scales on top (s.d., standard deviation). Lower end of voltage scale bars, –45 mV. Top right, normalized GCaMP impulse response. **e**, Two-dimensional histogram of measured vs. predicted GCaMP signals, color-coded according to the key on the right. *R*^2^=0.54±0.21 (mean±s.d.). **f**, Firing rate (30-ms moving average, black) aligned to GCaMP transient onset, defined as the zero crossing of the slope function of the normalized fluorescence trace (grey). **g**, Example whole-cell recordings from spontaneously spiking dFBNs. The coefficient of variation (CV) of the interspike interval (ISI) differs between tonic (<1), bursting (>1), and random (∼1) activity. Data are means ± s.e.m. For imaging details see Supplementary Table 1.

The regularity of GCaMP transients across the dataset in Fig. 1a raises the question of whether the entire dFBN population is engaged in synchronous oscillations or whether a subset of rhythmic cells dominates dendritic fluorescence, which represents a weighted sum of contributions from all dFBN processes in the imaged region. To distinguish these possibilities, we recorded simultaneous GCaMP signals from a median of 8 cell bodies (range: 3–12) in the same hemisphere (Fig. 1b, Supplementary Video 2). Time series of somatic fluorescence were, on average, uncorrelated (Pearson *r* = 0.05 ± 0.15, mean ± s.d). Different dFBNs concentrated variable amounts of power in the 0.2–2 Hz band, giving rise to a heavily skewed distribution (Fig. 1c, note the logarithmic scale). At the upper extreme of this continuum were neurons with pacemaker-like Ca^2+^ oscillations that resembled those in dendrites (Fig. 1b, c, Fig. S 1a–e). A handful of neurons could therefore account for much of the variance of the dendritic compound signal (mean *R*^2^=0.66, range: 0.46–0.81, Fig. S 1a–e), which in turn was tightly correlated with axonal fluorescence (Fig. S 1f–h).

To characterize the electrical events underlying GCaMP transients in slowly oscillating dFBNs, we performed simultaneous whole-cell patch-clamp and two-photon Ca^2+^-imaging experiments *in vivo*. The superimposition of voltage and GCaMP traces in Fig. 1d shows that Ca^2+^ transients coincided with large, sustained depolarizations generating high-frequency action potential bursts. These UP states alternated with DOWN states of somewhat longer average duration during which the neurons were hyperpolarized and stopped firing. For a quantitative analysis, we estimated the GCaMP impulse response from these recordings and used it successfully to predict fluorescence signals from the action potential record (Fig. 1d, e). A spike frequency histogram aligned to the onset of GCaMP transients peaked ∼120 ms before the maximal fluorescence change (Fig. 1f). The metronome-like alternations of UP and DOWN states that underpin the dendritic Ca^2+^ signal were merely one of several forms of temporally structured activity in the dFBN population: whole-cell recordings showed that some neurons released rhythmic bursts of action potentials at faster repetition rates than 0.5 Hz; others fired tonically at regular interspike intervals; still others produced repeating sequences of spikes and bursts (Fig. 1g). A few, but by no means all, of these activity patterns could also be discerned in the somatic Ca^2+^ traces of individual dFBNs (Fig. 1b, Fig. S 1a).

### A half-center oscillator generates SWA

Rhythmic neural activity is typically a product of cell-intrinsic oscillators, such as hyperpolarization-activated cation channels^8^, or oscillatory circuits, which abound in the neural control of movement^9-12^ but have also been proposed to regulate sleep cycle time^13,14^, if not sleep itself. In a prototypical oscillatory circuit, exemplified by the two-neuron swimming system of *Clione*^10^ or the myriad central pattern generators that populate invertebrate ganglia or the vertebrate brain stem, mutual inhibition between two half-centers controlling opponent sets of muscles produces self-perpetuating cyclic movement^9-12^.

Analyses of dFBN activity, along with connectomic evidence^15,16^, indicate that two mutually inhibitory half-centers are responsible for the generation of SWA. Simultaneous bilateral imaging showed that Ca^2+^ transients in dFBNs of the left and right hemispheres alternated out of phase (Fig. 2a, Supplementary Video 1), so that the occurrence of a fluorescence maximum in one hemisphere was associated, on average, with a minimum in the other (Fig. 2b, c). In keeping with the idea that direct inhibition from the contralateral hemisphere is the source of the negative fluorescence deflection, the GCaMP signal dipped deeper the higher was the mean SWA power in both hemispheres (Fig. 2d). As expected for two neuronal populations synchronized in antiphase, interhemispheric cross-correlations returned minima at a lag of 0 s and a period of ∼2 s, corresponding to 0.5 Hz (Fig. 2e, Fig. S 2a–c).

**Figure 2.**
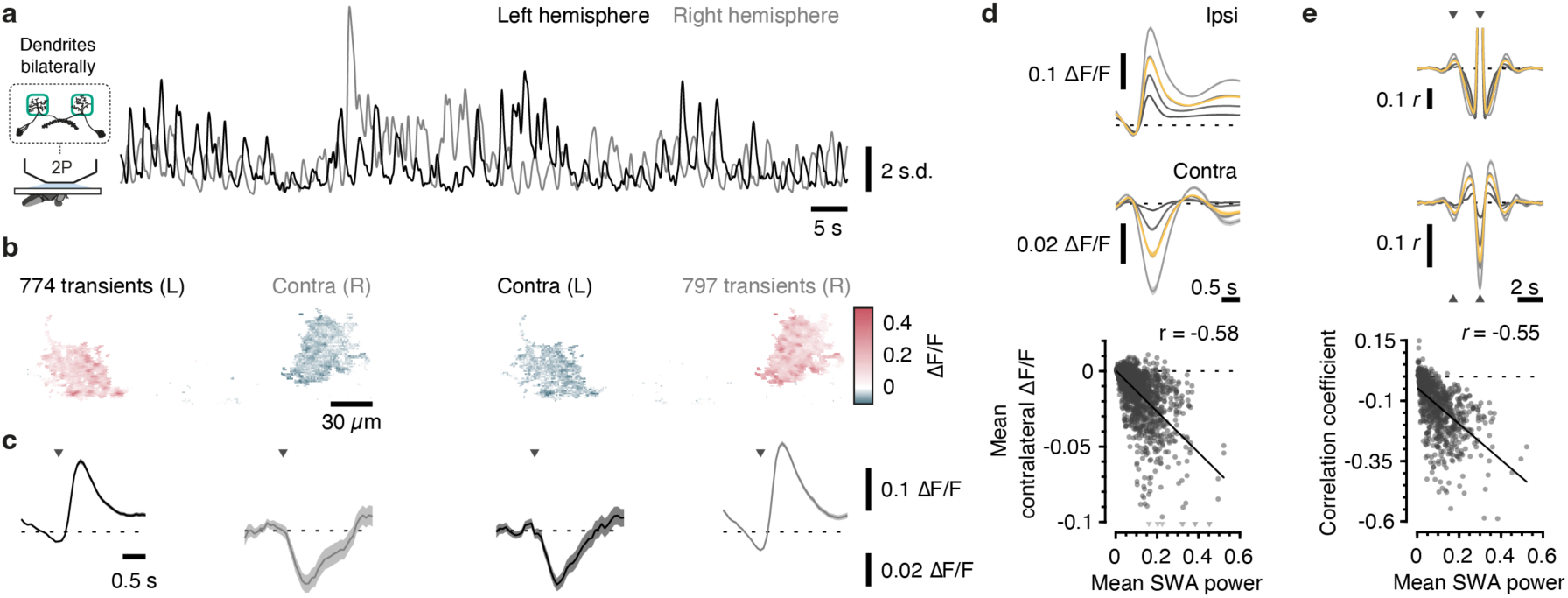
dFBNs form a half-center oscillator. **a**, Simultaneous imaging of dFBN dendrites in both hemispheres. The example shows 100 s of z-score-normalized GCaMP traces spanning ∼22 minutes. **b**, **c**, Ipsi-(red) and contralateral (blue) z-score-normalized fluorescence changes (**b**) and GCaMP traces (**c**) during transients in the left (L) and right (R) hemispheres (same fly as in **a**). Images in **b** depict mean ΔF/F in 10 frames (∼700 ms) after transient onset vs. the preceding 15 frames; the images are thresholded for display and pseudocolored according to the key on the right. Ipsi- and contralateral GCaMP traces in **c** are plotted at different ΔF/F scales; arrowheads mark the onset of ipsilateral transients; dashed horizontal lines indicate ΔF/F = 0. **d**, **e**, Transient-aligned average ipsi-(top) and contralateral (center) GCaMP traces (**d**) and their auto- and cross-correlograms (**e**) in the full dataset (*n*=284 flies; 790 ROIs). Shades of grey from dark to light show increasing quartiles of SWA power in both hemispheres; yellow traces represent population averages. Bottom, linear regression of mean contralateral ΔF/F (**d**) and interhemispheric correlation coefficients (**e**) vs. SWA power in both hemispheres. Nine data points exceeding the y-axis limit are plotted as triangles at the bottom of the graph in **d**; the correlation coefficient is based on the actual values. Shades of grey from dark to light show increasing quartiles of SWA power; yellow traces represent population averages. Correlograms are clipped near autocorrelation peaks; arrowheads indicate one period. Data are means ± s.e.m. For imaging details see Supplementary Table 1.

The symmetry and strength of connections among dFBNs in the hemibrain^15^ and FlyWire^16^ connectomes meets the anatomical requirements of a half-center oscillator (Fig. S 3). Almost perfectly symmetric weighted connectivity matrices (Fig. S 3a, f), each exhibiting a correlation coefficient of 0.92 or 0.81, respectively, with its transpose and a weight distribution indistinguishable from an undirected graph (Fig. S 3b, g), link the members of the population. Although dFBN interconnectivity is generally dense (full dFBN network: 0.66–0.73; dFBNs within layer 6: 0.94– 0.96; dFBNs within layer 7: 0.96–0.97) and reciprocal, regardless of synapse number, both connectomes favour connections within layers and between hemispheres and contain subsets of layer 6-projecting dFBNs^15^ with exceptionally strong connections with their contralateral counterparts (Fig. S 3a–j). We tentatively equate these neurons with dFBNs displaying a slow rhythm (Fig. 1c) but stress that a high degree of connection symmetry and reciprocity characterizes the dFBN population as a whole (Fig. S 3k, l).

Single-cell transcriptomic data reported in a companion paper^3^ identify glutamate as the fast-acting transmitter of dFBNs, in agreement with existing electrophysiological evidence that transmission from dFBNs (like glutamate release at many central synapses of *Drosophila*^17^) is inhibitory^18^. We confirmed this transmitter assignment anatomically and functionally. The subtraction of glutamatergic neurons from the *R23E10-GAL4* pattern (*R23E10-GAL4* ∖ *VGlut-GAL80*) retained two pairs of GABAergic cells in the suboesophageal zone and a handful of cholinergic neurons in the ventral nerve cord but excluded all dFBNs (Fig. 3a, b, Supplementary Video 3); conversely, the intersection of the *R23E10-GAL4* pattern with glutamatergic neurons (*R23E10 ∩ VGlut-GAL4*, created by reconstituting GAL4 from hemidrivers *R23E10-DBD* and *VGlut-p65AD*) highlighted dFBNs in the brain but lacked VNC-SP neurons^19^ of the nerve cord (Fig. 3c, Fig. S 4, Supplementary Video 4). dFBNs coexpressing the optogenetic actuator CsChrimson and the glutamate sensor iGluSnFR responded to light pulses with stimulus-locked fluorescence transients in their axonal fields, verifying synaptic glutamate release (Fig. 3d), whereas the acetylcholine sensors iAChSnFR and GRAB_ACh_ showed no evidence of cholinergic (co)transmission (Fig. 3d). Although some dFBNs transcribe genes encoding cholinergic machinery^3,20^, the translation of these mRNAs thus appears to be suppressed^21^.

**Figure 3.**
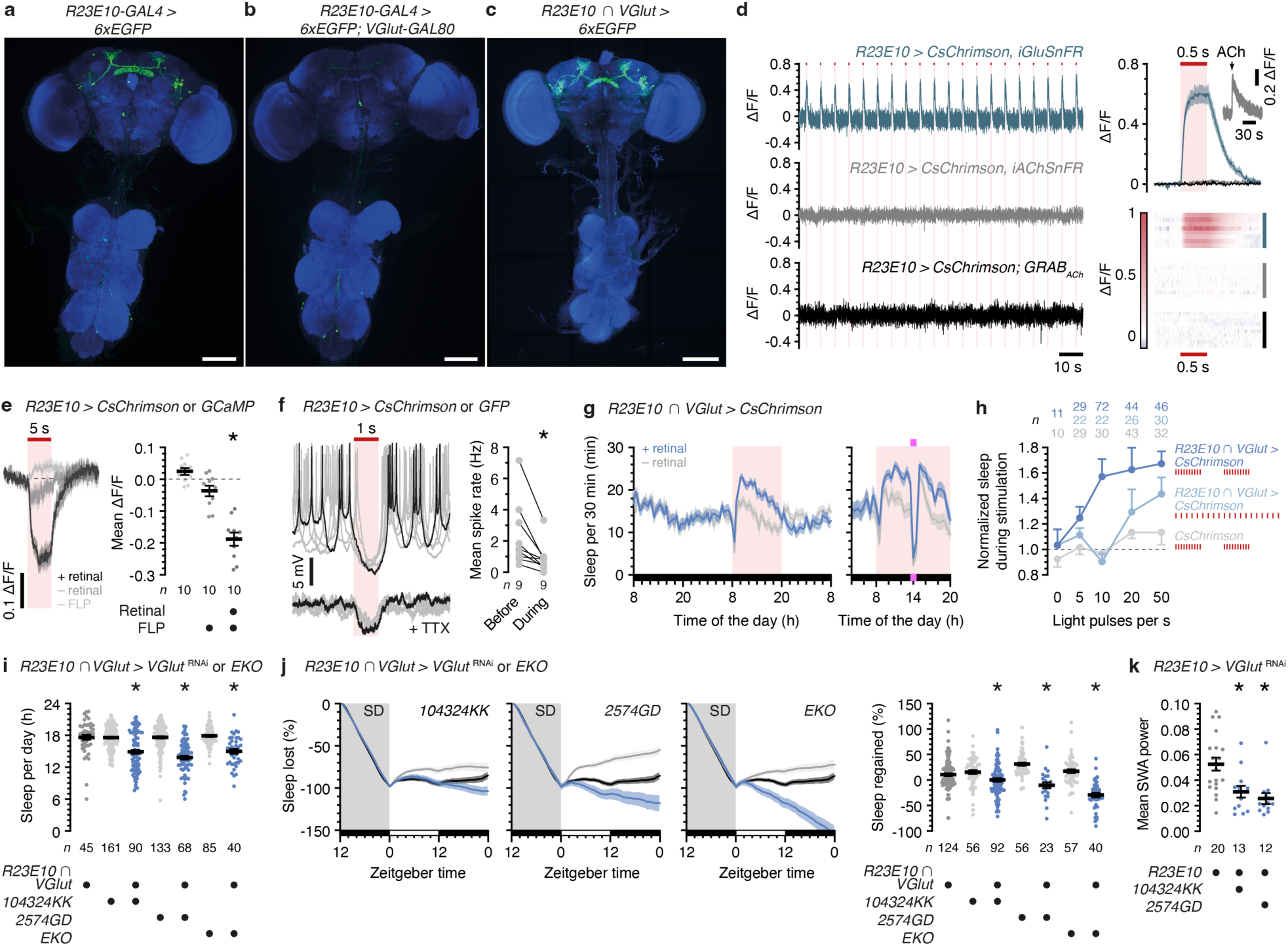
dFBNs promote sleep via glutamate. **a**, **b**, Maximum-intensity projections of the CNS of flies expressing 6xEGFP under the control of *R23E10-GAL4*, in the absence (**a**) or presence (**b**) of the transcriptional repressor VGlut-GAL80. **c**, Maximum-intensity projection of the CNS of a fly expressing 6xEGFP under the control of hemidrivers *R23E10-DBD* and *VGlut-p65AD* (*R23E10 ∩ VGlut-GAL4*). Green, 6xEGFP; blue, BRP. Scale bars, 100 μm. **d**, Optogenetic stimulation of dFBNs coexpressing CsChrimson and iGluSnFR (blue), iAChSnFR (grey), or GRAB_ACh_ (black) (*n*=8, 10, and 10, respectively). Left, background-corrected example traces during 20 light pulses (vertical bars, 500 ms). Top right, average stimulus-aligned, normalized fluorescence transients (inset, iAChSnFR after pressure ejection of acetylcholine). Bottom right, stimulus-aligned averages of individual flies, color-coded according to the key on the left. **e**, **f**, Imaging (**e**) and electrophysiological (**f**) demonstration of inhibition in dFBN mosaics expressing CsChrimson or a green fluorophore (GCaMP in **e**, mCD8::GFP in **f**) after FLP-mediated recombination. Somatic GCaMP fluorescence of CsChrimson-negative dFBNs declines during optogenetic stimulation of CsChrimson-positive dFBNs in the presence of retinal (**e**, *P*<0.0001, ANOVA). Optogenetic stimulation of CsChrimson-positive dFBNs hyperpolarizes CsChrimson-negative dFBNs (overlaid voltage traces of six consecutive trials, one trial in black) and suppresses spiking (**f**, *P*=0.0039, Wilcoxon test). Inter-dFBN inhibition persists in the presence of 1 µM tetrodotoxin (TTX). **g**, Sleep profiles (left) of flies expressing CsChrimson under the control of *R23E10 ∩ VGlut-GAL4*, with or without retinal (*n*=44 and 43, respectively), before, during, and after optogenetic replay of SWA (20 light pulses s^-1^ in 500-ms bursts) (retinal effect: *P*=0.0026, time × retinal interaction: *P*<0.0001, two-way repeated-measures ANOVA). (Ultra)violet illumination causes awakening during optogenetic stimulation (right, *n=*19 and 24 flies with and without retinal, respectively). **h**. Optogenetically induced sleep (normalized to non-retinal-fed controls) relative to carriers of an undriven *CsChrimson* transgene (genotype effect: *P*<0.0001, two-way ANOVA), as a function of the frequency and temporal structure of the optical pulse train (effect of number of light pulses s^-1^: *P*=0.0223, effect of temporal structure: *P*=0.0149, two-way ANOVA). **i**, **j**, *R23E10 ∩ VGlut-GAL4*-driven expression of *VGlut*^RNAi^ or EKO reduces daily sleep (**i**, *P*<0.0001, Dunn’s test after Kruskal-Wallis ANOVA) and the time courses and percentages of sleep regained after deprivation (SD) (**j**, left panels, genotype effects: *P*≤0.0087, time x genotype interactions: *P*<0.0001, two-way repeated-measures ANOVA; right panel, *P*≤0.0278, Dunn’s test after Kruskal-Wallis ANOVA). Same colors indicate genotypes throughout. **k**, *R23E10-GAL4*-driven expression of *VGlut*^RNAi^ reduces average SWA power (*P*=0.0005, Kruskal-Wallis ANOVA). Data are means ± s.e.m.; *n*, number of flies (**e, h**–**k**) or cells (**f**); asterisks, significant differences (*P*<0.05) in planned pairwise comparisons. For imaging details see Supplementary Table 1. For statistical details see Supplementary Table 2.

For a demonstration that dFBNs themselves are targets of dFBN-mediated inhibition, we divided the dFBN ensemble randomly, via stochastic recombination of GAL4-responsive FLP-out constructs^22^, into two disjoint groups containing either a green fluorophore (GCaMP for imaging; alternatively mCD8::GFP for targeted whole-cell electrophysiology) or CsChrimson::tdTomato (Fig. S 4h). Optical stimulation of the CsChrimson-expressing set resulted in reductions of GCaMP fluorescence in the complementary set of dFBNs in imaging experiments (Fig. 3e), or in hyperpolarization and the suppression of spiking in current-clamp recordings from GFP-positive (and therefore CsChrimson-negative) neurons (Fig. 3f). The addition of tetrodotoxin eliminated action potentials but not inhibitory postsynaptic potentials (Fig. 3f), which persist when transmission from directly connected terminals is triggered by light^22^. dFBN-to-dFBN communication is therefore monosynaptic.

The dense interconnectivity of dFBNs (Fig. S 3) explains the prevalence of mutual inhibition in mosaics (Fig. 3e, f) but complicates attempts to control the activity of the population. Hard, unrelenting, bilateral depolarization of dFBNs establishes a tug-of-war between the externally imposed excitation and the powerful all-to-all inhibition it elicits, with sometimes unforeseen consequences. Failures to observe increases in sleep after presumed (but unverified) activation of dFBNs led to a claim that the neurons are ineffective in promoting sleep^23^, when it was in fact the stimulation protocol that proved ineffective: pulsed illumination, which avoids a lasting cross-inhibitory stalemate, readily induced sleep in flies expressing CsChrimson under the control of *R23E10 ∩ VGlut-GAL4* exclusively in dFBNs, even though neurons of both hemispheres were inevitably driven in synchrony rather than their natural antiphase (Fig. 3g). Coincident exposures to (ultra)violet light cancelled the sleep-promoting effect of optogenetic stimulation and led to awakening (Fig. 3g). As in the regulation of sleep quality by clock neurons^24^ and the control of cortical SWA by the midline thalamus^25^, the temporal structure of the optical stimulus train mattered: increases in sleep quantity followed the simulated convergence of alternating 500-ms action potential bursts from the left and right hemispheres in the axonal target layers of dFBNs (which we approximated by compressing 5, 10, 20, or 50 light pulses into one half of a 500:500 ms duty cycle; Fig. 3g, h) but were lacking or lessened when the same number of pulses were spread evenly across time, mimicking tonic activity (Fig. 3h). The higher the intra-burst stimulation frequency was (a simulated increase in SWA power), the stronger was the sleep-promoting effect up to a saturation frequency of ∼40 Hz (Fig. 3h), approximating the peak intra-burst firing rate in our recordings of spontaneous dFBN activity (Fig. 1d).

To add evidence of necessity to this demonstration of sufficiency, we selectively incapacitated dFBNs by RNA-mediated interference (RNAi) with the expression of VGlut in the entire *R23E10-GAL4* territory, of which dFBNs are the sole glutamatergic members (Fig. S 4a–c), or by antagonizing glutamate release or membrane depolarization with the help of the intersectional driver *R23E10 ∩ VGlut-GAL4* (Fig. 3c, Fig. S 4c). Without exception (Fig. 3i, j, Fig. S 5), the depletion of VGlut from dFBNs or the introduction of the low voltage-activated potassium channel EKO reduced baseline and rebound sleep after mechanical sleep deprivation (using either of two methods; Fig. 3i, j, Fig. S 5a–d). Sleep losses occurred without potentially confounding waking hyperactivity (Fig. S 5a, c) or innervation defects that might have followed impaired glutamatergic transmission (Fig. S 6a), and they persisted when *teashirt-GAL80* eliminated *R23E10-GAL4*-driven *VGlut*^RNAi^ transgene expression in the ventral nerve cord (Fig. S 4e, Extened Data Fig. 6b) or a *tubulin-GAL80*^ts^-mediated block was relieved only after eclosion (Fig. S 6c), ruling out a developmental origin. Consistent with a critical role of reciprocal inhibition in antiphase oscillations^9-12^, interference with glutamatergic transmission also disrupted SWA (Fig. 3k, Fig. S 6d).

Experiments with *CsChrimson* and *VGlut*^RNAi^ transgenes under the control of a second split-GAL4 driver^20^ (*R23E10 ∩ R84C10-GAL4*), which, like *R23E10 ∩ VGlut-GAL4*, lacks expression in VNC-SP neurons of the ventral nerve cord (Fig. S 4f), confirmed that the output of the dFBN ensemble is somnogenic (Fig. S 5e) and required for the homeostatic regulation of sleep (Fig. S 5f, g).

### SWA encodes sleep pressure

The sleep deficits following the dissolution of SWA hint, and the sleep gains due to SWA replay over tonic stimulation strongly suggest, that slow waves have an instructive role in the translation of sleep pressure into sleep (Fig. 3g–k, Fig. S 5, Fig. S 6b–d). The electroencephalogram (EEG) during mammalian non-rapid eye movement (NREM) sleep^7,26^ is characterized by SWA that reflects synchronous transitions of cortical neurons between depolarized UP and hyperpolarized DOWN states^27^, not unlike those of rhythmic dFBNs (Fig. 1d). Cortical SWA is a classical EEG marker of sleep pressure: SWA levels at sleep onset increase after prolonged wakefulness and decrease after sleep^28^. To search for analogous modulations of dFBN activity, we compared SWA in flies that had been mechanically sleep-deprived for different spans of time with rested controls. Overnight sleep deprivation for >6 h (again using two different methods to similar effect) increased the average peak amplitude of GCaMP transients and the mean SWA power (Fig. 4a–d; Fig. S 2d, e) but left other frequency bands unchanged (Fig. 4c). As flies recovered, Ca^2+^ transients and SWA power decayed back to baseline over the course of 12–24 hours (Fig. 4e, Fig. S 2f). Following an unperturbed night of rest, in contrast, SWA was at its minimum in the morning (zeitgeber time 0–4), rose naturally during the day to an evening peak (zeitgeber time 12), and declined again during the night (Fig. 4e).

**Figure 4.**
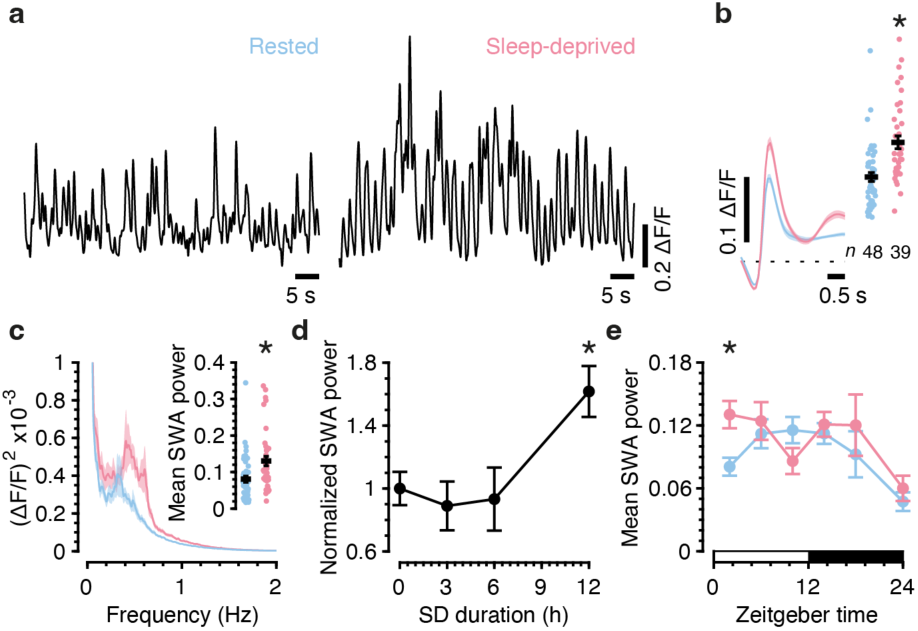
dFBN slow-wave activity encodes sleep need. **a**, Example GCaMP traces of dFBN dendrites at zeitgeber time (ZT) 0–4, after unperturbed rest (blue) or sleep deprivation between zeitgeber times 12 and 24 (red). **b**, **c**, Sleep deprivation increases the average GCaMP transient (**b**, left), its peak amplitude (**b**, right, *P*<0.0001, Mann-Whitney test), and SWA power (**c**, *P*=0.0004, Mann-Whitney test). **d**, SWA power (normalized to unperturbed controls) at ZT 0–4 as a function of the duration of prior sleep deprivation (*P*=0.0001, Kruskal-Wallis ANOVA). *n* at SD durations of 0, 3, 6, and 12 h: 48, 14, 12, 39 (same data at 0- and 12-h time points as in **b**, **c**). **e**, Mean SWA power during the course of a day in rested (blue) and sleep-deprived flies (*P*=0.0012, Kruskal-Wallis ANOVA). *n* of sleep-deprived and rested flies, respectively, at the given ZT were 0–4: 48, 39; 4–8: 32, 14; 8–12: 25, 14; 12–16: 31, 15; 16–23: 13, 9; 23–4: 9, 11. Data are means ± s.e.m.; *n*, number of flies; asterisks, significant differences (*P*<0.05) in planned pairwise comparisons. Same dataset as in Fig. 1a, Fig. 2, and Fig. S 2. For imaging details see Supplementary Table 1. For statistical details see Supplementary Table 2.

Previous reports of sleep-related rhythms^29,30^, and in particular observations of SWA in parts of the central complex^31,32^ (R5 neurons and dFBNs), noted but did not explore possible parallels with NREM sleep because SWA was characterized *ex vivo* and could therefore not be linked to a behavioral read-out of vigilance state. It is thus unclear if the occurrence of dFBN oscillations defines a slow-wave sleep stage as in mammals or if SWA tracks variations in sleep pressure throughout the sleep–wake cycle. To resolve this point, we examined if dFBN oscillations persisted during a behaviorally verified state of arousal. Head-fixed flies expressing either the dopamine sensor GRAB_DA2m_ or GCaMP in dFBNs were placed on a spherical treadmill and roused with a pulse of infrared laser light focused on their abdomina (Fig. 5a). In line with the notion that dopaminergic projections to the fan-shaped body mediate arousal by inhibiting dFBNs^6,33,34^, activating the laser for 2 s elicited a movement bout along with dopamine release onto the neurons’ dendrites (but not axons) (Fig. 5b). Like spontaneous spiking during dendritic applications of exogenous dopamine or optogenetic stimulation of dopaminergic neurons^6^, Ca^2+^ oscillations collapsed briefly during a short heat stimulus but resumed thereafter, following a powerful postinhibitory rebound (Fig. 5c–e), while pharmacological doses of dopamine produced pauses lasting many seconds^6^ (Fig. S 7).

**Figure 5.**
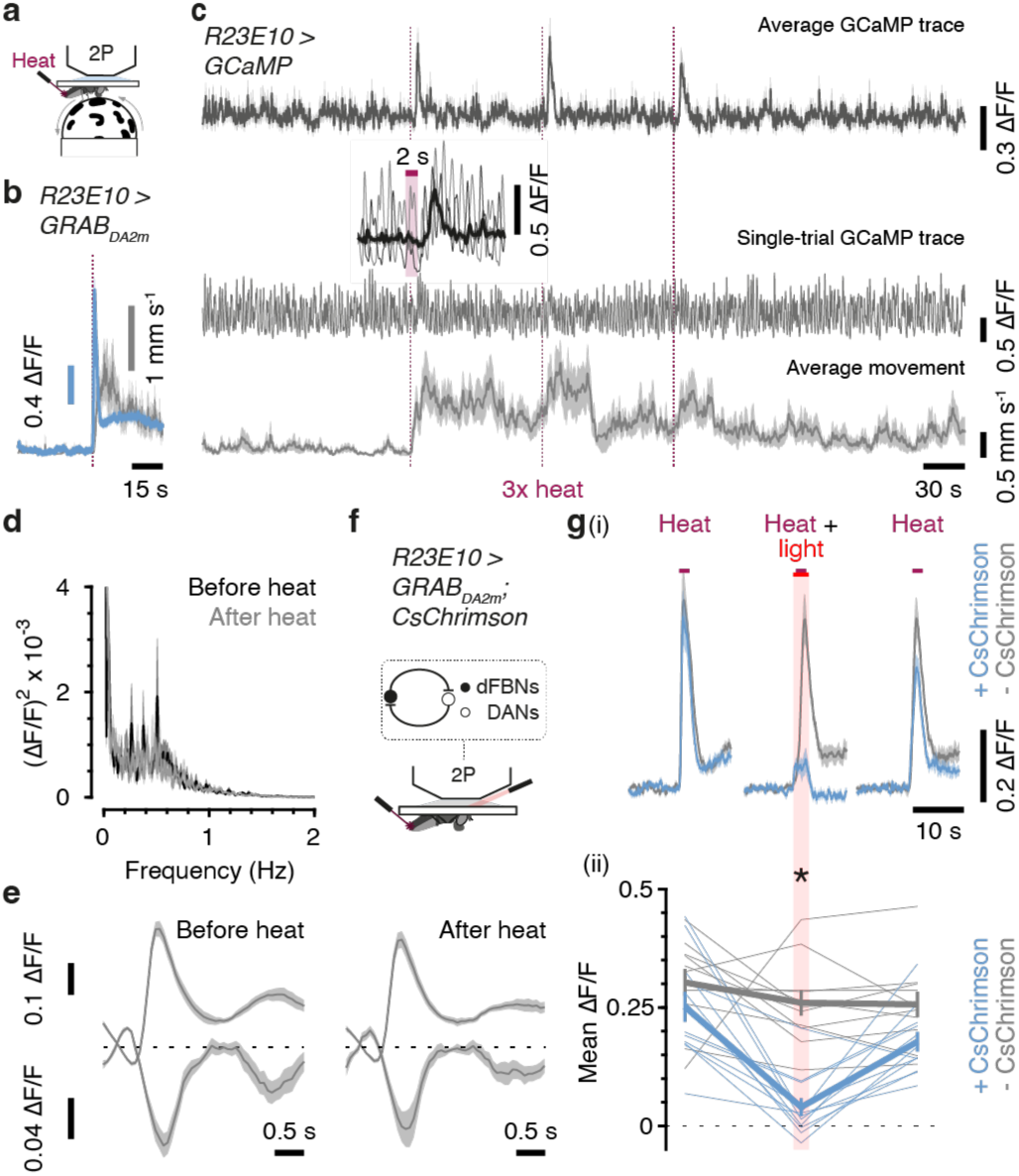
Slow-wave activity persists during arousal. **a**, Movement tracking and imaging during the application of arousing heat. **b**, Heat (dashed carmine line) stimulates dopamine release onto dFBN dendrites expressing GRAB_DA2m_ (blue) and locomotor bouts (grey) (*n*=9 flies). **c**, Dendritic GCaMP and locomotor traces before, during, and after three heat applications (dashed carmine lines). Top, cancellation of SWA in the average GCaMP trace (*n*=11 flies) reveals rebound responses aligned to the offset of heat. Lower end of scale bar, ΔF/F = 0. Center, GCaMP trace of a single trial. Lower end of scale bar, ΔF/F = –0.2. Segments of GCaMP traces in both hemispheres during one heat application and corresponding segment of the average trace (black) are shown at expanded x- and y-scales on top. Bottom, average locomotor speed. Lower end of scale bar, 0 mm s^-1^. **d**, **e**, Power spectra (**d**) and transient-aligned average ipsi-(top) and contralateral (bottom) GCaMP traces (**e**) in 8 flies where both hemispheres were imaged, during 140-s windows before and after the application of heat. **f**, Optogenetic stimulation of dFBNs during the application of heat probes for an inhibitory flip-flop arrangement of dFBNs and arousing dopaminergic neurons (DANs). **g**, Light during heat application inhibits dopamine release onto CsChrimson-positive (blue, *n*=12 flies) but not CsChrimson-negative dFBNs (grey, *n*=11 flies). Top (i), example traces of three applications of heat (spaced 30 minutes apart) to the same flies. Traces were smoothed with a 5-element (∼300 ms) moving-average filter for display. Bottom (ii), dFBN stimulation reduces mean GRAB_DA2m_ fluorescence during heat application (heat × CsChrimson interaction: *P*=0.0017, two-way repeated-measures ANOVA). Thin lines, individual flies; thick lines, population averages. Data are means ± s.e.m.; asterisk, significant difference (*P*<0.05) in a planned pairwise comparison. For imaging details see Supplementary Table 1. For statistical details see Supplementary Table 2.

Coincident optogenetic activation of dFBNs reduced the amplitude of heat-evoked dopamine transients (Fig. 5f, g), indicating that dFBNs and dopaminergic neurons antagonize each other in an arrangement reminiscent of the inhibitory flip-flop between sleep- and wake-promoting neurons in the mammalian hypothalamus^35^.

### Glutamate release adjusts to sleep pressure

In contrast to SWA in mammals, which can be recorded across the entire cortical surface during NREM sleep^7^, dFBN slow waves are products of a local circuit with features of a central pattern generator. Despite their apparent simplicity, pattern generators are notorious for the sensitivity of their dynamics to modulatory changes in biophysical parameters^36-38^. dFBNs experience many such changes as sleep pressure builds: they steepen their current–spike frequency functions^2^, modulate voltage-gated and leak potassium conductances in opposite directions^6^, and receive heightened (indirect) excitatory drive from R5 neurons^39^. Single-cell transcriptomic data reported in a companion study^3^ suggest that these changes, which all enhance activity, are counterbalanced by a weakening of efferent dFBN synapses. Sleep deprivation was found selectively to downregulate gene products with roles in synapse assembly, active zone maintenance, synaptic vesicle release, and presynaptic homeostatic plasticity^3,40^. Prominent among these differentially expressed genes was *bruchpilot* (*brp*)^3^, which encodes a structural component of active zones^41,42^. Tagging the endogenous BRP protein with a V5 peptide through dFBN-restricted genomic recombination^43^ allowed us to quantify its abundance as a function of sleep history. In a clear reflection of the transcriptional picture^3^, the intensity of BRP::V5 fluorescence (normalized to that of mCD8::GFP), an index of active zone size and a structural correlate of synaptic strength^44^, decreased at dFBN synapses of sleep-deprived flies (Fig. 6a). This is an unusual adjustment: waking experience, whether enforced by mechanical agitation or enriched by social interaction, is widely thought to lead to synaptic potentiation^45^, and axon terminal numbers or BRP levels indeed increase in many parts of the sleep-deprived *Drosophila* brain^46,47^, including virtually all specific neuron types that have been examined, such as R5 neurons^39^ (Fig. S 8a), Kenyon cells^48^, and clock neurons^46^. The anti-cyclical depression of dFBN synapses after sleep loss thus underlines, like their anti-cyclical energy metabolism does^3^, their special status with respect to sleep.

**Figure 6.**
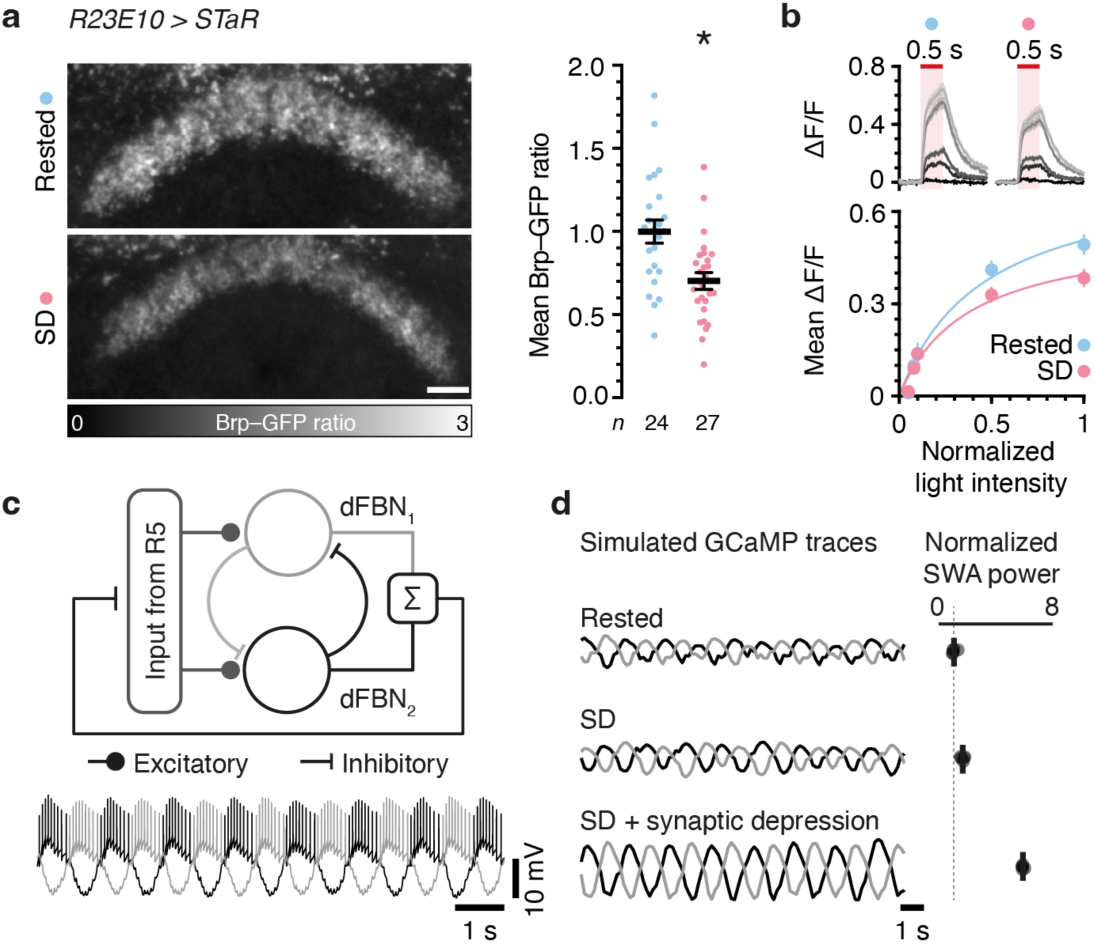
Efferent dFBN synapses depress after sleep deprivation. **a**, Summed-intensity projections of V5-tagged endogenous BRP in dFBN axons coexpressing mCD8::GFP. Emission ratios are intensity-coded according to the key at the bottom and decrease after sleep deprivation (SD) (*P*=0.0009, *t*-test). Scale bar, 10 μm. **b**, Sleep deprivation reduces glutamate release elicited by optogenetic stimulation of dFBNs (light intensity × sleep history interaction: *P*=0.0002, two-way repeated-measures ANOVA). Top, iGluSnFR traces (*n*=11 rested and 12 SD flies). Shades of grey from dark to light show increasing light intensities normalized to peak (25 mW cm^-2^). Bottom, mean iGluSnFR ΔF/F during illumination in rested (blue) and sleep-deprived flies (red) as functions of light intensity. Data are fitted by saturation hyperbolas. **c**, Minimal model of two dFBNs with reciprocal inhibitory connections in a simplified feedback circuit with a pool of excitatory Poisson units. Top, model schematic. Bottom, simulated voltage traces of dFBNs color-coded as on top. **d**, Simulated GCaMP traces (left) and mean SWA power (right) at baseline and after sleep deprivation, in the absence and presence of plastic feedback connections with Poisson units (*n*=5 simulations). Data are means ± s.e.m.; asterisk, significant difference (*P*<0.05) in a planned pairwise comparison. For imaging details see Supplementary Table 1. For statistical details see Supplementary Table 2.

For a functional confirmation under activity-normalized conditions that sleep deprivation attenuated synaptic transmission from dFBNs, we measured the optically evoked axonal release of glutamate from dFBNs co-expressing CsChrimson and iGluSnFR. Saturation hyperbolas described the dependence of transmitter secretion on light intensity irrespective of sleep history, but the iGluSnFR signal saturated at lower levels in sleep-deprived than in rested flies (Fig. 6b). The significance of presynaptic depression, which we have now demonstrated at the transcriptional^3^, structural, and functional levels, for the sleep need-dependent modulation of SWA becomes clear in a simple model where two dFBNs are embedded in recurrent circuitry^18,39^ and connected across the midline (Fig. 6c, Fig. S 8b). These direct, symmetric, and inhibitory connections are necessary (and must be sufficiently strong) to produce rhythmic, pacemaker-like SWA. Empirically, individual dFBNs become more excitable as sleep pressure rises^2^; artificially enhancing this excitability promotes sleep^3-6^; SWA power grows with sleep drive (Fig. 4c, Fig. S 2e); and optogenetic replay of SWA induces sleep (Fig. 3g, h, Fig. S 5e). However, the network’s recurrent organization allows an increase in SWA following changes in intrinsic dFBN excitability and external excitatory drive only if a homeostatic weakening of dFBN synapses is part of the simulations (Fig. 6d); without it, dFBNs would cut off their own sources of excitation (Fig. S 8c). The two seemingly opposing forces observed in dFBNs after sleep deprivation—increased excitability and decreased synaptic output— may in reality work together to generate an oscillatory representation of sleep need.

## Discussion

Our work, along with previous studies^29-32^, has uncovered remarkable parallels and striking differences in the sleep-related neuronal dynamics of flies and mammals. Slow waves reflecting alternating UP and DOWN states are a common marker of sleep pressure, but the biophysical origin, spatial spread, and temporal persistence of these waves differ. A central pattern generator with rhythmic dFBNs at its core encodes sleep need in the amplitude of its periodic output, which oscillates robustly also in awake flies. It is this stability—and the considerable lengths dFBNs go to to maintain it at variable levels of sleep pressure—that poses the most intriguing questions.

dFBNs keep a record of sleep need in a biochemical memory where each unit of storage is a Shaker– Hyperkinetic channel whose cofactor oxidation state holds one bit of information^5^. Productive encounters with PUFA-derived carbonyls increase, whereas depolarization-controlled exchange reactions decrease, the fraction of channels with oxidized cofactors^5^. The fill level of the integrator thus depends on a dynamic equilibrium between oxidation and exchange. Regular spikes would tip this equilibrium toward exchange and thereby introduce a leak into the integrator, extending its time constant. We currently do not know how rapidly the Hyperkinetic pool reaches capacity, but the addition of a spike-triggered leak offers flexibility to adjust the time to saturation. With a range of spiking patterns in the dFBN population, which is likely to include forms of coordinated activity that have remained undetected by Ca^2+^ imaging, the same mechanism could in principle produce a spectrum of integration times and the characteristic heavy-tailed distribution of sleep bout durations.

dFBNs have a demonstrated ability to switch reversibly between long-lasting electrically active ON and quiescent OFF states^6^. These binary states (which were discovered under conditions where transmembrane currents were clamped^6^) bracket the extremes of the dynamic range used by dFBNs to map largely cell-autonomous, sleep history-dependent metabolic variables, such as mitochondrial electron usage^3,4^ or levels of peroxidized lipids^5^, onto electrical signals that can be communicated to other cells. Under unclamped conditions, we now understand, this dynamic range is set so that an oscillatory representation of sleep pressure continuously flows to downstream structures, including during wakefulness. Only manipulations which forcibly block the discharge of sleep pressure appear to interrupt this periodic signal—at a price that is debited biochemically^3-5^ and repaid in heightened SWA power when inhibition is released. dFBNs fall silent during applications of exogenous dopamine^6^ and reduce their activity far below baseline during mechanical sleep deprivation^49^; conversely, artificial depolarization of dFBNs during enforced waking prevents the mitochondrial rearrangements that are a visible token of the associated metabolic cost^3^.

The utility of communicating an unbroken record of sleep pressure to the action-selection circuitry of the fan-shaped body^15^ is obvious: the likely need for future sleep is an important factor in choosing what to do now. However, if the same periodic signal also controls transitions between sleep–wake states, as our replay and interference experiments confirm, continuous variations in SWA power must be transformed into binary outcomes. This analog-to-digital conversion could take several forms. For example, dFBNs could target a dedicated sleep module^15^ in the central complex through strongly facilitating synapses that selectively transmit^50^ the high-frequency, large-amplitude somnogenic bursts we observe. Alternatively, because action selection is an inherently competitive winner-take-all process, it may be sufficient for dFBNs to alter the probability that inaction is favoured among competing options and let the internal architecture of the fan-shaped body^15^ secure the exclusivity of the choice.

## Methods

### Drosophila strains and culture

Flies were grown on media of cornmeal (62.5 g l^-1^), inactive yeast powder (25 g l^-1^), agar (6.75 g l^-1^), molasses (37.5 ml l^-1^), propionic acid (4.2 ml l^-1^), tegosept (1.4 g l^-1^), and ethanol (7 ml l^-1^) under a 12 h light:12 h dark cycle at 25 °C in ∼60% relative humidity, unless stated otherwise. To prevent the undesired activation of optogenetic actuators or the photoconversion of all-*trans* retinal by ambient light, flies expressing CsChrimson and their controls were reared and housed in constant darkness and transferred to food supplemented with 2 mM all-*trans* retinal (Molekula or Sigma) in DMSO, or to DMSO vehicle only, 2 days before the optical stimulation experiments, at an age of 1–3 days post eclosion. Carriers of the *hs-FLP* transgene^51^ were cultivated at 18 °C and placed on retinal-supplemented food for 5 days before optogenetic stimulation experiments; carriers of the *tub-GAL80^ts^*transgene and their controls were raised at 18°C.

Driver lines^52,53^ *R23E10-GAL4* and *R23E10*-*LexA*; the repressor lines *VGlut-3xGAL80*^54^, *teashirt-GAL80*^55^, and *tubulin-GAL80^ts^* (ref. ^56^), and the split-GAL4 hemidrivers^54,57-59^ *R23E10-DBD*, *VGlut-AD*, *R84C10-AD*, and *Gad1-AD* were used in the indicated combinations to target dFBNs or other constituents of the *R23E10-GAL4* pattern at fine genetic resolution; *R69F08-GAL4* labelled R5 ring neurons of the ellipsoid body^39,52^. Effector transgenes encoded fluorescent markers for neuroanatomy (*UAS-6xEGFP*^60^, *UAS-mCD8::GFP*^61^, *lexAop-GFP*^62^, *UAS-mCD4::tdTomato*^63^, *UAS-myrRFP*) or the labelling of synaptic active zones^43^ (*lexAop-GFP*;*UAS-STaR*); the Ca^2+^ sensor^64,65^ GCaMP6f; the glutamate^66^, acetylcholine^67,68^, or dopamine^69^ sensors iGluSnFR(A184V), GRAB_ACh_, iAChSnFr, or GRAB_DA2m_, respectively; the optogenetic actuator^70^ CsChrimson^71^; cassettes^22^ for the mutually exclusive expression of green fluorescence (CD8::GFP or GCaMP6f) or CsChrimson::tdTomato after the FLP-mediated excision^51^ of a transcriptional terminator; the low voltage-activated Shaker derivative^72^ EKO; or RNAi constructs for interference with the expression of *VGlut* (4 independent transgenes^73,74^).

### Sleep measurements, sleep deprivation, and optogenetic induction of sleep

Females aged 2–4 days were individually inserted into 65-mm glass tubes containing food reservoirs, loaded into the Trikinetics *Drosophila* Activity Monitor system, and housed under 12 h light:12 h dark conditions at 25 °C in ∼60% relative humidity. Flies were allowed to habituate for one day before sleep— classified^75,76^ as periods of inactivity lasting >5 minutes (Sleep and Circadian Analysis MATLAB Program^77^)—was averaged over two consecutive recording days (Fig. S 9a–d). Immobile flies (< 2 beam breaks per 24 h) were manually excluded. Flies carrying *tub-GAL80^ts^* transgenes and their controls were transferred to 31 °C for two days before sleep measurements, also at 31 °C, or kept at 18 °C throughout.

Our standard method of sleep deprivation used the sleep-nullifying apparatus^78^ (SNAP): a spring-loaded platform stacked with Trikinetics monitors was slowly tilted by an electric motor, released, and allowed to snap back to its original position. The mechanical cycles lasted 10 s and were repeated continuously. Alternatively, as noted, an Ohaus Vortex Mixer stacked with Trikinetics monitors produced horizontal circular motion stimuli with a radius of ∼1 cm at 25 Hz for 2 s; stimulation periods were randomly normally distributed within 20-s bins. Unless otherwise stated, sleep deprivation lasted for 12 h, beginning at zeitgeber time 12. The two methods produced comparable levels of sleep loss and rebound (Fig. 3j, Fig. S 5b, d, Fig. S 9a–d) without impairing post-deprivation locomotion (Fig. S 9e).

Rebound sleep was measured in the 24-h window after deprivation (Fig. S 9a–d). A cumulative sleep loss plot was calculated for each individual by comparing the percentage of sleep lost during overnight sleep deprivation to the immediately preceding unperturbed night. Individual sleep regained was quantified by normalizing the difference in sleep amount between the rebound and baseline days to baseline sleep. Only flies losing >95% of baseline sleep were included in the analysis.

In photostimulation experiments, female flies expressing CsChrimson in dFBNs were individually inserted into 65 mm glass tubes and loaded into a custom array^18^ of light-tight chambers with high-power 630-nm LEDs (Multicomp OSW-4388). The apparatus was operated in a temperature-controlled incubator (Sanyo MIR-154) at 25 °C. For movement tracking, the chambers were continuously illuminated from below using low-power infrared (850 nm) LEDs and imaged from above with a high-resolution CMOS camera (Thorlabs DCC1545M) equipped with a long-pass filter (Thorlabs, FL850-10) to reject stimulation light pulses, which lasted 3 ms and delivered ∼28 mW cm^-2^ of optical power. A virtual instrument written in LabVIEW (National Instruments) extracted real-time position data from video images. Periods of inactivity lasting ≥5 minutes were classified as sleep. If flies were found dead at the end of the two-day experiment, data from the 30-min time bin preceding the onset of continuous immobility (≥29.5 min per 30-min bin for ≥2 h until the end of the experiment) onward were excluded; if the period of continuous immobility began during or before optical stimulation, all data from that individual were discarded. The 29.5-min threshold was applied to avoid scoring rare instances of video tracker noise as movement.

To rouse flies during optogenetic stimulation of dFBNs, a 385-nm LED (M385L3, ThorLabs) controlled by a dimmable LED driver (LEDD1B, ThorLabs) delivered ∼1–3 mW cm^-2^ of optical power continuously for 3 minutes, followed by 3 minutes of darkness. Cycles lasted for 1 h, beginning at zeitgeber time 13.5.

### Two-photon imaging

Females aged 1–3 days were implanted with chronic imaging windows^79,80^ 2– 3 days before the experiment. Unless otherwise stated (Supplementary Table 1), surgical openings created by removing cuticle, adipose tissue, and trachea were sealed with a thin layer of translucent UV-curable epoxy glue (Norland NOA13825; Opticure 2000 UV lamp, Norland) in a custom chamber perfused with CO_2_ to avoid O_2_ inhibition of the curing process, which lasted <45 s. The adhesive stabilized the brain, had a refractive index matched to that of water, and lacked any detectable adverse effects—including effects on sleep—after flies had recovered from anaesthesia (Fig. S 10a, b). Flies were housed singly for 2–3 days after the procedure, cold-anaesthetized, head-fixed to a custom mount with eicosane (Sigma), allowed to recover for ≥20 minutes, and imaged on a Movable Objective Microscope with resonant scanners (MOM, Sutter Instruments) controlled through ScanImage software (Vidrio Technologies). For sleep deprivation before imaging, experimental flies and their controls were placed into Trikinetics monitors 1–2 days after the implantation of the window.

Only healthy flies in which the adhesive effectively suppressed brain movements were imaged. Fluorophores were excited by a Mai Tai DeepSee Ti:sapphire laser (Spectra Physics model eHP DS; center wavelength of 930 nm). A Pockels cell (302RM, Conoptics) kept the power at the specimen <15 mW (ThorLabs PM100D power meter console with ThorLabs S370C sensor head). Emitted photons were collected by a 20×/1.0 NA water immersion objective (W-Plan-Apochromat, Zeiss), split into green and red channels by a dichromatic mirror (Chroma 565dcxr) and bandpass filters (Chroma ET525/70m-2p and Chroma ET605/70m, respectively), and detected by GaAsP photomultiplier tubes (H10770PA-40 SEL, Hamamatsu Photonics). If simultaneous optogenetic stimulation was performed, an alternative bandpass filter (Semrock BrightLine FF01-520/60-25) was present in the green emission path. Photocurrents were passed through high-speed amplifiers (HCA-4M-500K-C, Laser Components) and integrators (BLP-21.4+, Mini-Circuits) to maximize the signal-to-noise ratio. The objective was mounted on a MIPOS piezo actuator (PiezoSystemJena) controlled through ScanImage. Supplementary Table 1 lists the number of imaging planes, their axial distances, numbers of pixels, acquisition rates, and additional parameters for each experiment.

For optogenetic stimulation, a 625-nm LED (M625L3, ThorLabs) controlled by a dimmable LED driver (LEDD1B, ThorLabs) delivered ∼0.5–25 mW cm^-2^ of optical power through a bandpass filter (Semrock BrightLine FF01-647/57-25) to the head of the fly. The voltage steps controlling the LED driver were recorded in a separate imaging channel for post-hoc alignment. Imaging of iGluSnFR, iAChSnFR, and GRAB_ACh_ during optogenetic stimulation of dFBNs in Fig. 3d was performed in the presence of 1 µM tetrodotoxin (TTX).

To deliver arousing stimuli, an 808-nm laser diode (Thorlabs L808P500MM) was mounted on a temperature-controlled heat sink (ThorLabs TCDLM9 with ThorLabs TED200C controller) and aimed at the abdomen of a fly standing or walking on a 6-mm polystyrene ball supported by a stream of air (0.3 l min^-1^). The diode was restricted to a maximal output of 50 mW by a ThorLabs LDC210C laser diode controller. Rotations of the spherical treadmill were recorded by a GuppyPro camera equipped with a magnification lens and a Semrock FF01-647/57-50 bandpass filter at a mean acquisition rate of 22.80 Hz. The treadmill was dimly illuminated by a red LED through a bandpass filter (Semrock BrightLine FF01-647/57-25). Forward and turning velocities were computed by a custom MATLAB script (Jan Kropf, C.B.T., G.M., in preparation). Movement and imaging traces were aligned post hoc using simultaneously recorded voltage steps as time stamps.

For focal dopamine or acetylcholine applications, patch pipettes (∼10 MΩ) containing 10 mM dopamine or 30 mM acetylcholine in extracellular solution were positioned near the dendritic (in the case of dopamine) or axonal (in the case of acetylcholine) arborizations of dFBNs^6^. A TTL-controlled voltage step triggered the delivery of transmitter by a PDES-02DX pneumatic drug ejection system (npi electronic GmbH). Voltage steps were recorded in ScanImage for post-hoc alignment. Movement artefacts were controlled for by titrating the ejection pressure to 10–25 kPa and recording mCD4::tdTomato fluorescence in a separate imaging channel. In the case of acetylcholine, the pulse duration was 40 ms, close to the minimal valve switching time.

Functional imaging data were analysed using custom code in MATLAB. Rectangular regions of interest (ROIs) containing dFBN processes or irregularly shaped ROIs following the contours of dFBN somata or axons were manually drawn, along with a background ROI positioned on a non-fluorescent brain area, and time series of mean fluorescence intensities were extracted from two-photon image stacks. Following the subtraction of the average intensity of the background time series, ΔF/F curves were computed as ΔF_t_/F^0^_t_ = (F_t_ – F^0^_t_)/ F^0^_t_, where F_t_ is the raw fluorescence intensity in frame t. For event-aligned analyses (i.e., GCaMP transient onset, optogenetic stimulation, arousing heat), F^0^_t_ was the mean fluorescence in a given time window before the event. To correct for motion during pharmacological dopamine applications, normalized changes in green-to-red ratios (ΔR/R) were computed as ΔR_t_/R_0_ = (R_t_ – R_0_)/ R_0_, where R_t_ is the element-wise ratio between background-subtracted GCaMP and mCD4::tdTomato signals and R_0_ is the mean ratio 40 s before dopamine application.

For measurements of spontaneous dFBN activity, the time-varying baseline fluorescence F^0^_t_ was obtained as the 10^th^ percentile of a symmetric sliding window of F_t_, spanning 501 frames (∼34 s). The resulting ΔF/F traces were smoothed with an 8-element Gaussian sliding window. GCaMP transients were detected as peaks in the low-pass filtered (15-element Gaussian sliding window) time derivative of the ΔF/F trace; the amplitudes of transients were defined as the maximum within the 15 frames following the maximal rise of ΔF/F. Interhemispheric cross-correlations were computed on the normalized time derivative of the traces, which eliminates slow trends in the signal but captures the time course of dFBN bursts (Fig. 1f). For the calculation of coherence spectra, time series were windowed into 500-element long segments overlapping by 50%.

The blinded extraction of ROIs required some exclusions because of low signal-to-noise ratios. ROIs were excluded automatically from further analysis if their background-subtracted mean GCaMP fluorescence failed to exceed the standard deviation of a comparable background signal by 20-fold (raw traces) or 45-fold (smoothed traces). The baseline fluorescence of most ROIs easily passed these thresholds, with mean GCaMP fluorescence exceeding the standard deviation of background noise by factors of 92 (raw traces) or 193 (smoothed traces). In addition, where several focal planes of the same hemisphere were imaged, signals were screened for multiple representations of the same cell(s) by computing pairwise correlations between ROIs at different focal depths. Only the ROI showing the largest fluorescence intensity was retained if the correlation coefficient exceeded 0.85 for dendritic ROIs and 0.5 for somatic ROIs; in the latter case we confirmed by visual inspection that redundant ROIs covered the same somata. In GRAB_DA2m_ experiments, the ROI displaying the highest response to the first heat shock was selected automatically for further analysis.

To relate GCaMP signals recorded simultaneously in somata and dendrites (Fig. S1a–e) or dendrites and axons (Fig. S1f–h), linear regression models using the somatic or dendritic ΔF/F traces as predictor variables of the dendritic or axonal traces, respectively, were fit in MATLAB. Model performance was evaluated by 5-fold cross-validation.

Because Ca^2+^ diffuses into dFBN somata through the long, thin necks of dendrites and axons, somatic GCaMP signals are temporally low-pass filtered versions of signals recorded in the neuropil. This made an exhaustive characterization and classification of activity patterns of the dFBN ensemble (whose members could be resolved only in cell body images) impossible. Our analysis therefore focused on the distinction of pacemaker-like activity, which was readily detectable, from all other forms of activity. ΔF/F traces of individual somata were used to compute the peak signal power between 0.2 and 2 Hz; a prominent periodicity in the autocorrelogram with a wide amplitude swing during the first period was diagnostic of a slow rhythm.

### Electrophysiology

Female flies were head-fixed to a custom mount with eicosane (Sigma) or soft thermoplastic wax (Agar Scientific). A surgical window was cut into the cuticle; adipose tissue, trachea, and the perineural sheath were removed; and the brain was superfused with carbogenated extracellular solution (95% O_2_ – 5% CO_2,_ pH 7.3, 275 mOsm) containing (in mM) 103 NaCl, 3 KCl, 5 TES, 8 trehalose, 10 glucose, 7 sucrose, 26 NaHCO_3_, 1 NaH_2_PO_4_, 1.5 CaCl_2_, 4 MgCl_2_. The GFP- or GCaMP- and mCD4::tdTomato-labelled somata of dFBNs were visualized using 20×/1.0 NA (W-Plan-Apochromat, Zeiss), 40×/0.8 NA, or 60×/1.0 NA water immersion objectives (LUMPLFLN40XW or LUMPLFLN60XW, Olympus) and targeted with borosilicate glass electrodes (8–11 MΩ) filled with internal solution (pH 7.3, 265 mOsM) containing (in mM): 10 HEPES, 140 potassium aspartate, 1 KCl, 4 MgATP, 0.5 Na_3_GTP, 1 EGTA. Signals were acquired at room temperature in current-clamp mode with a MultiClamp 700B amplifier (Molecular Devices), lowpass-filtered at 5–20 kHz, and sampled at 10–50 kHz using an Axon Digidata 1440 digitizer controlled through pCLAMP 10.5 (Molecular Devices) or a Power1401-3A data acquisition interface controlled through Signal (Cambridge Electronic Design Ltd.).

For simultaneous whole-cell patch-clamp recording and two-photon imaging, a TTL-gated voltage signal was recorded in pClamp and ScanImage for the post-hoc alignment of electrophysiology and imaging data. Simultaneously recorded membrane voltages were compared with GCaMP fluorescence time series at the soma (9 of 10 cells); in one cell, Ca^2+^ signals were imaged in dendrites. The ΔF/F curve was computed as ΔF_t_/F^0^_t_ = (F_t_ – F^0^_t_)/ F^0^_t_, where F_t_ is the raw fluorescence trace and the time-varying baseline fluorescence F^0^_t_ was obtained as a 300-element (∼5 s) moving average of F_t_. The imaging time series was z-score-normalized and linearly interpolated and patch-clamp recordings were down-sampled, so that both signals had the same temporal resolution of 2 kHz; the resulting traces were aligned to the TTL time stamp. The momentary firing rate (or peri-event time histogram, PETH) was computed in 100-ms bins and used to create a predictor matrix for the estimation of the GCaMP impulse response from the interpolated and aligned imaging trace, which was smoothed using a 150-ms moving average and then downsampled to the same frame rate as the predictor matrix (10 Hz). The resolution of 10 Hz was chosen to control zero inflation in the predictor matrix (i.e., the number of time bins without spikes in the PETH) and reduce the number of coefficient weights to be fitted to 20 for a 2-s long impulse response. Model performance was evaluated by estimating the individual GCaMP impulse responses of all 10 cells by linear regression and predicting the Ca^2+^ signal of each dFBN from its spiking record, using the normalized impulse response derived from the other 9 cells (leave-one-out cross-validation). Coefficients of determination (*R*^2^) were obtained from the squared maximum cross-correlations between normalized and predicted imaging data.

In studies of inter-dFBN connectivity, low basal expression of the *hsFLP* transgene, without additional heat shock, was sufficient to produce recombination events^22,51^ whose visible sign was a mosaic of green and red fluorescent dFBNs (Fig. S 4h). The presence of both dFBN populations was confirmed by live microscopy before each experiment. CsChrimson-expressing cells were stimulated with a 630-nm LED (Multicomp OSW-4388) controlled by a TTL-triggered dimmable LED driver (Recom RCD-24-0.70/W/X3) and focused on the fly’s head with a mounted 60-mm lens (Thorlabs). The light source delivered 11–80 mW cm^-2^ of optical power. Where indicated, 1 μM tetrodotoxin (TTX) was perfused into the bath to probe for monosynaptic connections^22,81^. Data were analysed with custom procedures, using the NeuroMatic package^82^ (http://neuromatic.thinkrandom.com) in Igor Pro (WaveMetrics).

### Confocal imaging

Single-housed females aged 6 days post eclosion were dissected at zeitgeber time 0. For the quantification of BRP::V5, experimental and control samples were processed in parallel, following *ad libitum* sleep or 12 h of sleep deprivation. Nervous systems were fixed for 20 min in 0.3% (v/v) Triton X-100 in PBS (PBST) with 4% (w/v) paraformaldehyde, washed five times with PBST, and incubated sequentially at 4 °C in blocking solution (10% goat serum in 0.3% PBST) overnight, with primary antibodies in blocking solution for 2–3 days, and with secondary antibodies in blocking solution for two days. The primary antibodies included mouse nc82 anti-BRP (1:10, Developmental Studies Hybridoma Bank), mouse anti-V5 (1:400, ThermoFisher), chicken anti-GFP (1:1000 or 1:500 for STaR, AbCam), and mouse anti-GFP (1:50 for GCaMP, ThermoFisher); the secondary antibodies were goat anti-Mouse Alexa Fluor 488 (1:1000, ThermoFisher), goat anti-Chicken Alexa Fluor 488 (1:1000 or 1:500 for STaR, ThermoFisher), and goat anti-Mouse Alexa Fluor 633 (1:500, ThermoFisher). The samples were washed five times with PBST before and after the addition of secondary antibodies, mounted in Vectashield, and imaged on a Leica TCS SP5 confocal microscope with an HCX IRAPO L 25×/0.95 water immersion objective. Only anatomically intact specimens from live flies (at the point of dissection) were analysed.

For evaluating possible anatomical anomalies after interference with the expression of VGlut in dFBNs, brains expressing *R23E10 ∩ VGlut-GAL4*-driven GFP and *VGlut^RNAi^* or *RFP* transgenes were counterstained for BRP, imaged, and analysed with a semi-automated custom script in Fiji. The BRP signal was used to define the dFB volume manually. GFP fluorescence was quantified in summed z-stacks containing this volume, following the subtraction of average background fluorescence in a nearby area. To estimate the density of dFBN projections into this volume (i.e., the innervation density), z-slices were passed through a Gaussian low-pass filter (0.5 σ) and manually thresholded to exclude isolated low-intensity pixels. GFP-positive volumes were identified and measured with the ‘3D Objects Counter’ function in Fiji (‘threshold’ = 10, ‘minimum puncta size’ = 100 voxels), summed, and expressed as a percentage of the dFB volume.

### Connectome analysis

The hemibrain:v1.2.1 connectome^83^ and the FlyWire female adult fly brain dataset FAFB v783 (refs. ^16,84,85^) were accessed in neuPrint+ (ref. ^86^) and Codex (https://codex.flywire.ai), respectively. The total number of synapses per connection was used as measure of connection strength. Connectivity matrices take into account cells assigned to *R23E10-GAL4* in the hemibrain^15^ and cell types FB6E1, FB6E3, FB6E4, and FB7E2 in the FlyWire^85^ connectomes; the same classification was used for comparisons between and within cell types.

Connectivity matrices were computed and analysed in MATLAB. Omitting all diagonal entries (the number of autapses, which was uniformly zero), an index of network density was obtained as the number of observed edges (inter-dFBN connections) divided by the number of all possible edges in the graph. The original directed graph was compared with an undirected graph, which was created by averaging the weighted connectivity matrix with its transpose. Instances of unidirectional connections formed by one synapse were assigned a mean of 1 in the undirected graph. Non-zero entries above the diagonal of the undirected connectivity matrices were binned to create the histograms of Fig. S 3b, g. To avoid potential confounds arising from the heavy-tailed weight distribution and high network density, the reciprocity of inter-dFBN connections was quantified as a function of connection strength. To do so, the weighted connectivity matrices were binarized with thresholds *s* ranging from 0 to 100, resulting in series of Boolean matrices *B*^(*s*)^. The reciprocity *r*(*s*) of connections with synapse number *n* > *s* was obtained as

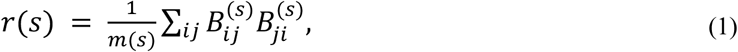

where *m*(*s*) is the total number of connections (non-zero entries) in *B*^(*s*)^. Connection symmetry (that is, the relative strength of reciprocal connections) was evaluated by comparing the weighted connectivity matrix *A* with its transpose *A^T^*. The Pearson correlation coefficient (omitting diagonal entries) of *A* and *A^T^* approaches 1 while the difference between *A* and *A^T^* approaches zero for a perfectly symmetric matrix. The scaled distance *a* from the zero matrix was therefore used as a measure of asymmetry:

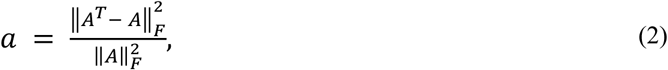

where 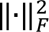 is the squared Frobenius norm (equivalent to the sum of squares of all entries) and 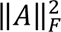 is used for normalization. For null model comparisons (*n*=100,000), all entries of *A* (and consequently *A^T^*) were randomly permuted before computing *a*. Values close to the mean of the respective null models are a sign of asymmetric connectivity (see, for example, MBONs in Fig. S 3l).

### Quantification and statistical analysis

Imaging and behavioral data were analysed in MATLAB and Prism 10 (GraphPad). All null hypothesis tests were two-sided. To control type I errors, *P*-values were adjusted to achieve a joint α of 0.05 at each level in a hypothesis hierarchy; multiplicity adjusted *P*-values are reported in cases of multiple comparisons at one level. Group means were compared by *t* test, one-way ANOVA, two-way repeated-measures ANOVA, or mixed-effects models, as stated, followed by planned pairwise analyses using Holm-Šidák’s multiple comparisons test. Repeated-measures ANOVA and mixed-effect models used the Geisser-Greenhouse correction. Where the assumption of normality was violated (as indicated by D’Agostino-Pearson test), group means were compared by Mann-Whitney test or Kruskal-Wallis ANOVA, followed by Dunn’s multiple comparisons test, as indicated.

The investigators were blind to sleep history, zeitgeber time, the inclusion or exclusion of dietary retinal, and/or genotype during the selection of ROIs and background regions in functional imaging experiments (Fig. 1a, Fig. 2, Fig. 3e, k, Fig. 4, Fig. 5g, Fig, 6b, Fig. S 1f–h, Fig. S 2, Fig. S 6d), the analysis of dFBN projections after interference with glutamatergic transmission (Fig. S 6a), and STaR experiments (Fig. 6a, Fig. S 8a) but not otherwise (Supplementary Table 1). Sample sizes in behavioral experiments (typically *n*=32 flies per genotype) were chosen to detect 2-h differences in daily sleep with a power of 0.9. All behavioral experiments were run at least three times, on different days and with different batches of flies. The figures show pooled data from all replicates.

### Modelling

We studied a model of two dFBNs (one in the left and one in the right hemisphere) with reciprocal inhibitory connections. Each dFBN was driven by 20 presynaptic units whose spike trains were inhomogeneous Poisson processes; the rates of these Poisson processes were subject to feedback modulation by the summed inhibitory output of both dFBNs. The architecture of the model is based on anatomical and functional evidence of recurrent connectivity between R5 neurons of the ellipsoid body (corresponding to the presynaptic units) and dFBNs^18,39^. R5 neurons provide indirect excitation to dFBNs^39^, whose inhibitory activity in turn diminishes (again via indirect connections^18^) the excitatory drive R5 neurons provide. Our minimal model recapitulates the basic feedback logic of the circuit but simplifies it by replacing indirect with direct connections and relying on a biphasic filter (rather than explicit conductances) to model hyperpolarization and hyperpolarization-induced escape.

Sleep loss-dependent changes were incorporated via three model parameters, as suggested by empirical data (Fig. S 5b). First, dFBNs in sleep-deprived flies increase their intrinsic excitability^2^. This was modelled as a shortening of the refractory period following a dFBN spike. Second, R5 neurons display higher firing rates and larger presynaptic active zones after sleep deprivation^39^ (Fig. S 5a). This was modelled as an increase in the rate of the Poisson units. Third, dFBN output synapses weaken after sleep loss (Fig. 6a, b). This was modelled as a decrease in the connection weight between dFBNs and the Poisson units.

A spike response model^87^ described the membrane potential *v_j_*(*t*) of each dFBN in discrete 1-ms time steps as a linear sum of a resting membrane potential (*v_rest_*) of –50 mV, excitatory postsynaptic potentials (EPSPs) due to input from *n* = 20 Poisson units, spike after-hyperpolarization, and the inhibitory postsynaptic potentials (IPSPs) and postinhibitory rebound elicited by the contralateral dFBN:

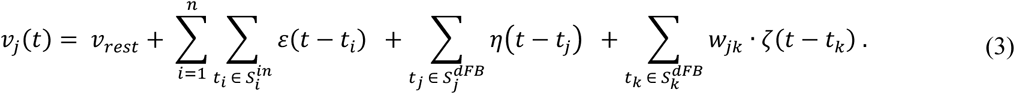

*S_i_^in^*, *S_j_^dFB^*, and *S_k_^dFB^* are sets of spike times (*t_i_*, *t_j_*, and *t_k_*) of the *i^th^* input unit, dFBN*_j_*, and the contralateral dFBN*_k_*.

EPSPs had a rise time constant *τ*_1_ of 7 ms and a decay time constant *τ*_2_ of 150 ms; their amplitude *ε* was scaled to a peak of 0.75 mV:

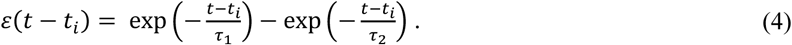

The post-spike hyperpolarization was defined as

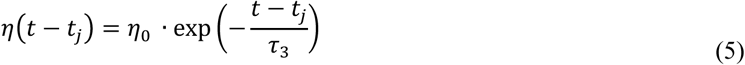

with *τ*_3_ = 300 ms and *ƞ*_0_ = −6 mV.

The time course of an IPSP and its postinhibitory rebound was encapsulated by a biphasic filter

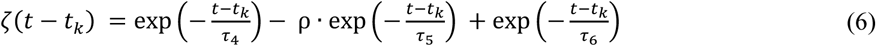

with *τ*_4_ = 450 ms, *τ*_5_ = 225 ms, *τ*_6_ = 56 ms, and ρ = 2. The filter was normalized and weighted by the amplitude *w_jk_* = *w_kj_*_’_ = 8 mV for each contralateral dFBN spike (Eq. 3).

Spike initiation by dFBNs was deterministic. If a time-varying voltage threshold *ϑ*(*t*) was crossed, an action potential was initiated (modelled as the addition of a 20-mV impulse to the current threshold value). To incorporate refractoriness, this deterministic threshold was a function of spiking history

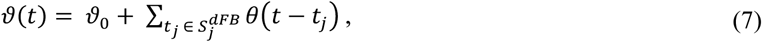

where the baseline spiking threshold *ϑ*_0_ was –45 mV and

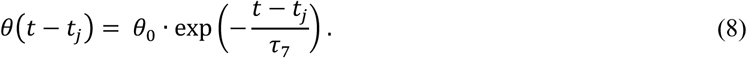

The decay time constant *τ*_7_ was 30 ms in rested and 15 ms in sleep-deprived conditions, respectively, to account for variation in the intrinsic excitability of dFBNs^2^ (Fig. S 8b). *θ*_0_ was 400 mV.

The spike trains of 20 input units per dFBN were modelled as inhomogeneous Poisson processes with a basal rate *ω*_0_ of 30 in rested and 60 in sleep-deprived flies (Fig. S 8b), matching empirical firing rate increases in R5 neurons^39^. Inhibitory feedback from the summed output of both dFBNs reduced the momentary Poisson rate *ω*(*t*) according to

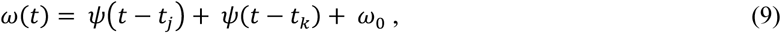

where

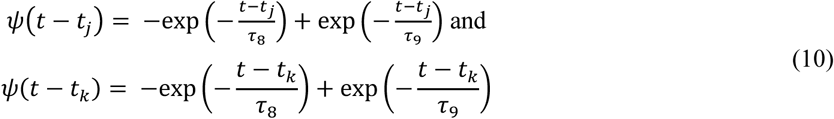

with *τ*_8_ = 7 ms and *τ*_9_= 1000 ms. The amplitude of *ψ* was normalized and scaled by a variable output weight reflecting the observed presynaptic plastic changes in dFBNs; sleep deprivation was modelled as a 50% reduction in the postsynaptic response amplitude (Fig. S 8b). *ω*(*t*) was restricted to positive values.

Ca^2+^ imaging experiments were simulated by convolving the momentary firing rates of model dFBNs in 100-ms bins with the GCaMP impulse response function estimated from simultaneous patch-clamp recordings. Data were z-score normalized.

## Acknowledgments

We thank C. Velasco for contributions to an analysis of dFBN types, G. Bertuzzi and D. Pimentel for preliminary imaging experiments, E. Agnes for discussions of the spike-response model, and R. Davis, K. Golic, H. Keshishian, Y.-N. Jan, V. Jayaraman, D. Kim, T. Lee, Y. Li, L. Luo, L. Looger, G. Rubin, R. Stowers, J. Truman, B. White, L. Zipursky, the Bloomington Stock Center, the Vienna *Drosophila* Resource Center, and the Transgenic RNAi Project (TRiP) for flies. This work was supported by grants from the European Research Council (832467) and Wellcome (209235/Z/17/Z and 106988/Z/15/Z) to G.M. P.S.H. was an Erwin Schrödinger Fellow of the Austrian Science Fund (J 4060); R.S. held a Wellcome Four-Year PhD Studentship in Basic Science (215200/Z/19/Z).

## Author contributions

P.S.H. performed and analysed all functional imaging studies, developed the dFBN model, analysed connectomic data with R.S., and wrote an initial manuscript draft; R.S. performed and analysed all neuroanatomical and behavioral and some imaging experiments; E.V. and H.O.R. obtained electrophysiological recordings; C.B.T. developed instrumentation; R.B. generated fly strains. P.S.H., R.S., and G.M. designed the study, interpreted the results, and prepared the manuscript. G.M. devised and directed the research and wrote the final version of the paper. Correspondence and requests for materials should be addressed to G.M.

---

**Figure S 1.**
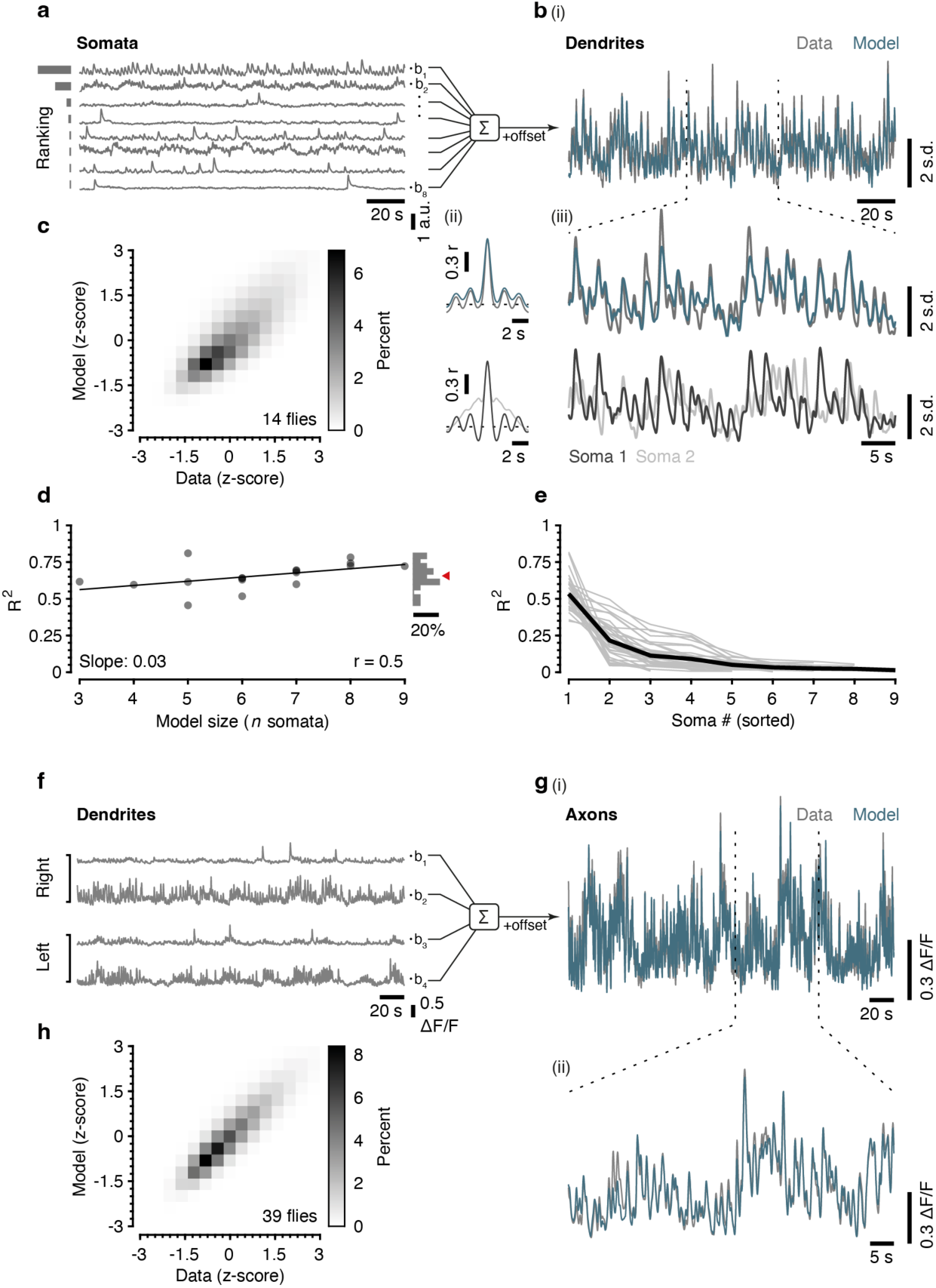
Simultaneous imaging of dFBN somata and dendrites, or dendrites and axons. **a**, **b**, Example GCaMP traces of dFBN somata (**a**) and dendrites (**b**). A linear model using the somatic signals as predictor variables (blue) accurately describes the dendritic GCaMP trace (**b**(i), grey), which is dominated by dFBNs with a slow rhythm (ii) and shown at an expanded time scale below (iii). Cells in **a** are ranked by the proportion of variation in dendritic fluorescence predicted by somatic fluorescence (see **e**). **c**, Two-dimensional histogram of measured vs. predicted dendritic GCaMP signals (*n*=14 flies, 17 separately recorded hemispheres, 35 dendritic ROIs). Data in the histogram were z-score-normalized for display but not for fitting and are color-coded according to the key on the right. **d**, Model performance (*R^2^*) as a function of the number of simultaneously imaged dFBNs. The vertical histogram summarizes the dataset; red arrowhead, mean *R^2^*. **e**, Proportion of variation in dendritic fluorescence predicted by somatic fluorescence, in decreasing order of *R^2^*. **f**, **g**, Example GCaMP traces of dFBN dendrites (**f**) and axons (**g**). A linear model using the dendritic signals as predictor variables (blue) accurately describes the axonal GCaMP trace (**g**(i), grey, shown at an expanded time scale below (ii)). **h**, Two-dimensional histogram of measured vs. predicted axonal GCaMP signals. Data in the histogram were z-score-normalized for display but not for fitting and are color-coded according to the key on the right. For imaging details see Supplementary Table 1.

**Figure S 2.**
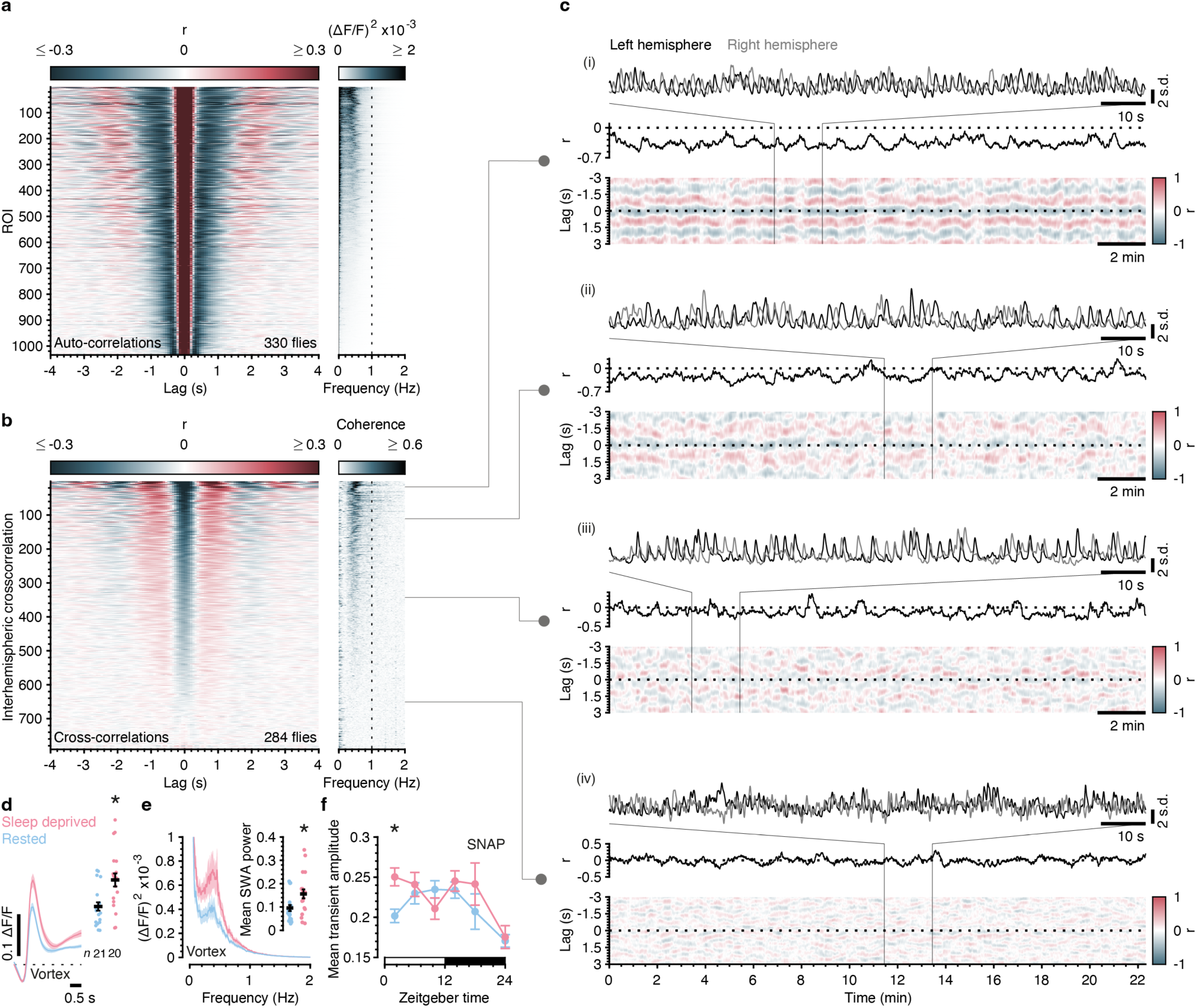
dFBN ensemble dynamics as a function of sleep need and zeitgeber time. **a**, Auto-correlograms (left) and power spectra (right) of GCaMP traces in all dendritic regions of interest (ROIs), sorted (in descending order) according to SWA power and color-coded according to the respective keys on top. **b**, Cross-correlograms (left) and coherence spectra (right) of dendritic GCaMP traces during imaging of both hemispheres, sorted (in descending order) by the average interhemispheric anticorrelation at a lag of 0±300 ms and color-coded according to the respective keys on top. **c**, Time-varying cross-correlograms (sliding 20-s windows, bottom panels) and correlation coefficients (center panels) of dendritic GCaMP traces in the left and right hemispheres at four levels of coherence. Top panels, left and right dendritic GCaMP traces at an expanded time scale. **d**, **e**, Sleep deprivation by vortex increases the average GCaMP transient (**d**, left), its peak amplitude (**d**, right, *P*=0.0013, *t*-test), and SWA power (**e**, *P*=0.0122, *t*-test). **f**, Mean amplitude of GCaMP transients during the course of a day in rested (blue) and sleep-deprived (red) flies (SNAP method; *P*<0.0001, Kruskal-Wallis ANOVA). Same dataset as in Fig. 4e. Data are means ± s.e.m.; asterisks, significant differences (*P*<0.05) in planned pairwise comparisons. For imaging details see Supplementary Table 1. For statistical details see Supplementary Table 3.

**Figure S 3.**
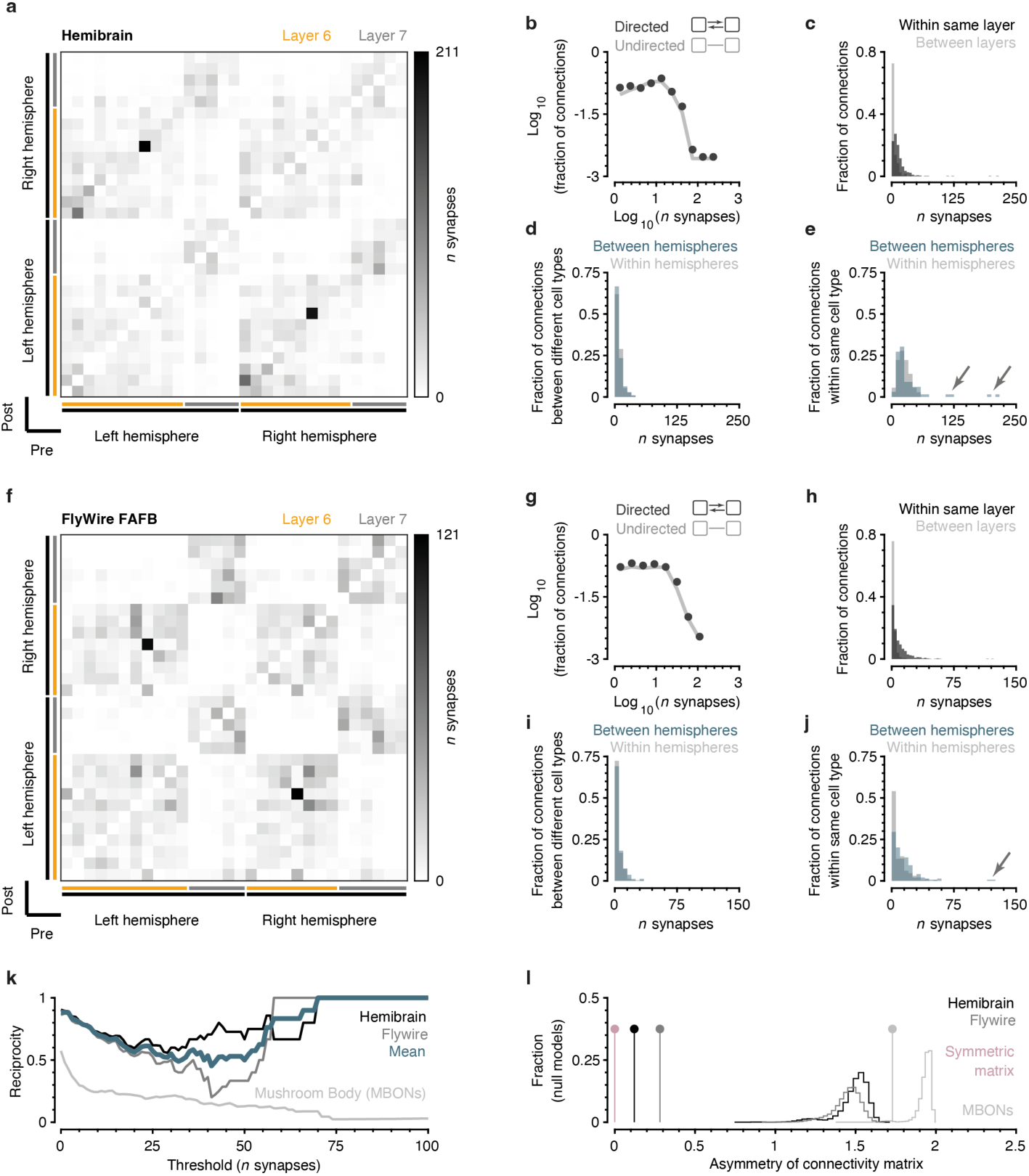
Connectomic support for a half-center oscillator. **a**, Connectivity matrix of dFBNs in the hemibrain:v1.2.1 dataset (*n*=31 dFBNs of both hemispheres, projecting to layers 6 and 7). Presynaptic neurons in rows, postsynaptic neurons in columns. **b**, Number of synapses per inter-dFBN connection. Conversion of directed (dark grey) to undirected edges (light grey) leaves the histogram unchanged. **c**, Inter-dFBN connections within (black) and between (grey) layers. dFBNs interconnect preferentially within their respective layers (*P*<0.0001, Mann-Whitney test). **d**, **e**, Inter-(green) and intra-hemispheric connections (light grey) between different (**d**) and the same (**e**) hemibrain dFBN types. The average number of synapses between the same dFBN types (29.96) exceeds the average number of synapses between different types (7.62) (*P*<0.0001, Mann-Whitney test). **f**, Connectivity matrix of dFBNs in the FlyWire FAFB v783 dataset (*n*=30 dFBNs of both hemispheres, projecting to layers 6 and 7). Presynaptic neurons in rows, postsynaptic neurons in columns. **g**, Number of synapses per inter-dFBN connection. Conversion of directed (dark grey) to undirected edges (light grey) leaves the histogram unchanged. **h**, Inter-dFBN connections within (black) and between (grey) layers. dFBNs interconnect preferentially within their respective layers (*P*<0.0001, Mann-Whitney test). **i**, **j**, Inter-(green) and intra-hemispheric connections (light grey) between different (**i**) and the same (**j**) hemibrain dFBN types. The average number of synapses between the same dFBN types (10.54) exceeds the average number of synapses between different types (4.20) (*P*<0.0001, Mann-Whitney test). **k**, Reciprocity of inter-dFBN connections as a function of synapse number. Black, grey, and blue traces are hemibrain, FlyWire, and the average of both, respectively. Light grey trace, connectivity of mushroom body output neurons (MBONs; FlyWire dataset). **l**, Asymmetry of inter-dFBN connectivity matrices of hemibrain and FlyWire connectomes, compared to their null models (black and grey stems and histograms, respectively). Values close to the mean of the respective null models are a sign of asymmetric connectivity. Light grey stem and histogram, MBONs. Red stem, symmetric matrix. For statistical details see Supplementary Table 3.

**Figure S 4.**
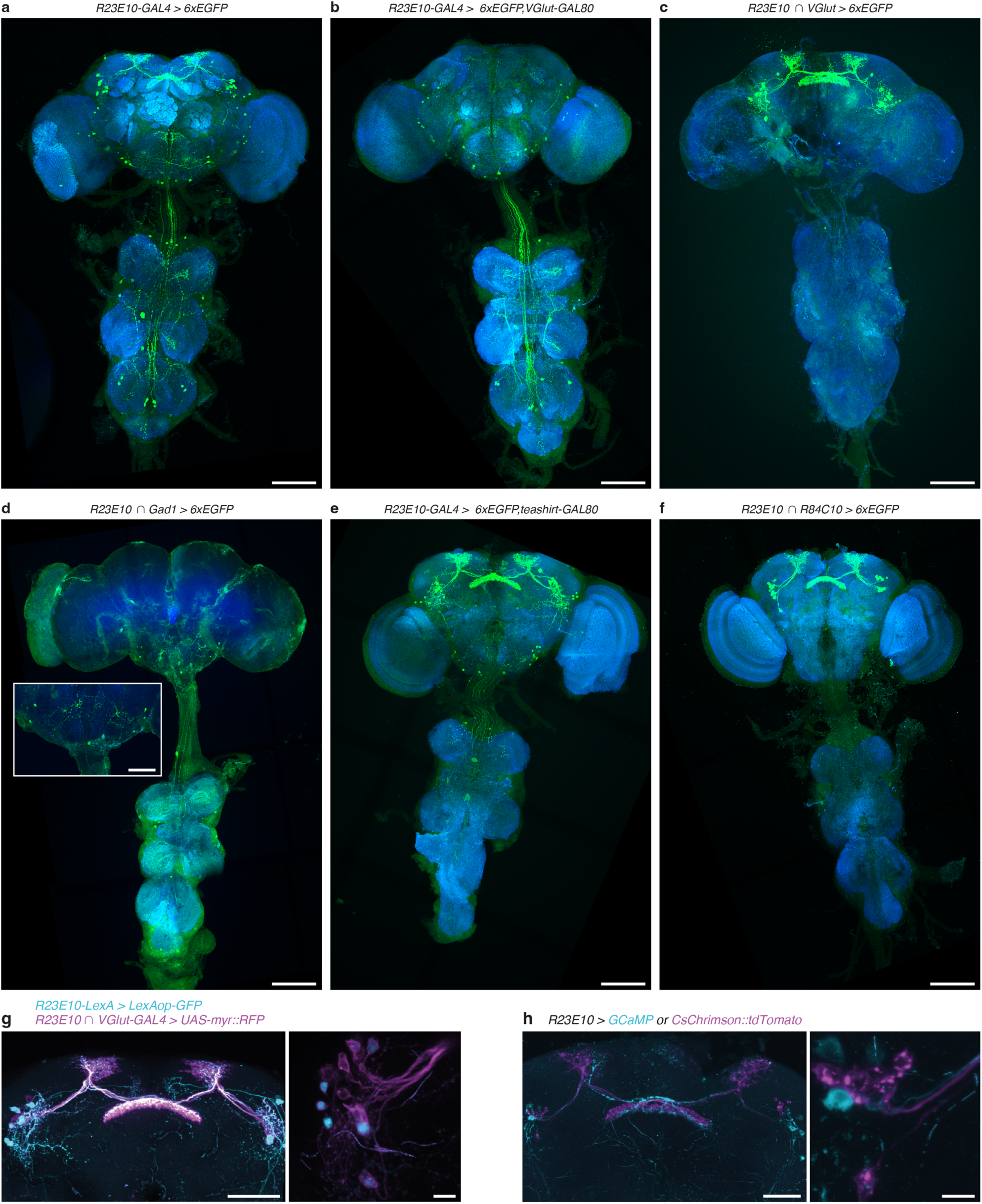
Genetic resolution of dFBNs. **a**–**f**, Saturated maximum-intensity projections of the CNS of flies expressing 6xEGFP under the control of *R23E10-GAL4*, in the absence (**a**) or presence of the transcriptional repressors VGlut-GAL80 (**b**) or teashirt-GAL80 (**e**), or under the control of hemidrivers *R23E10-DBD* and *VGlut-p65AD* (**c,** *R23E10* ∩ *VGlut-GAL4*), *R23E10-DBD* and *Gad1-p65AD* (**d**), or *R23E10-DBD* and *R84C10-p65AD* (**f**, *R23E10* ∩ *R84C10-GAL4)*. The inset in **d** shows a magnified view of the suboesophageal zone. Green, 6xEGFP; blue, BRP. Scale bars, 100 μm (inset in **d**, 50 μm)**. g**, Maximum-intensity projections of fly brains expressing GFP under the control of *R23E10-LexA* and myr::RFP under the control of *R23E10* ∩ *VGlut-GAL4* (left, whole midbrain; right, dFBN somata). *R23E10* ∩ *VGlut-GAL4* captures all dFBNs labelled by *R23E10-LexA*. Additional myr::RFP-positive dFBNs are included in the *R23E10-GAL4* pattern but missing from *R23E10-LexA*. Scale bars, 50 μm (left) and 10 μm (right). **h**, Maximum-intensity projections of fly brains expressing CsChrimson::tdTomato (magenta) or GCaMP (cyan) in a mutually exclusive fashion after FLP-mediated recombination (left, whole midbrain; right, dFBN somata). Scale bars, 50 μm (left) and 10 μm (right).

**Figure S 5.**
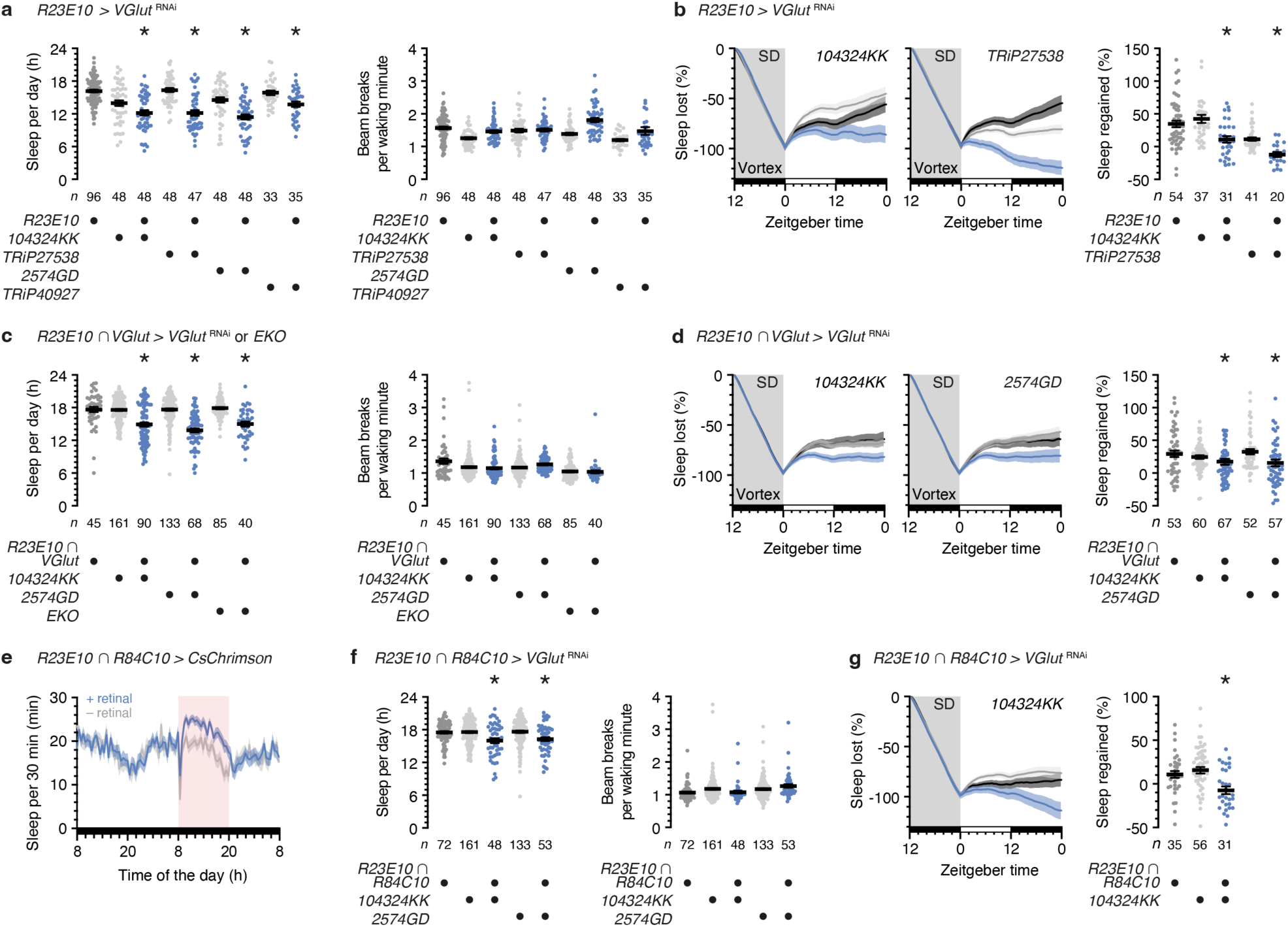
Different strategies for interfering with the function of dFBNs have consistent effects on sleep. **a**, **b**, *R23E10-GAL4*-driven expression of *VGlut*^RNAi^ reduces daily sleep (**a**, left, *P*≤0.0077, Holm-Šídák test after ANOVA) and the time courses and percentages of sleep regained after deprivation by vortex (SD) (**b**, left panels, genotype effects: *P*≤0.0176, time × genotype interactions: *P*<0.0001, two-way repeated-measures ANOVA; right panel, *P*≤0.0109, Dunn’s test after Kruskal-Wallis ANOVA) without altering waking locomotor activity (**a**, right, *P*≥0.1601 relative to ≥1 parental control, Dunn‘s test after Kruskal-Wallis ANOVA). **c**, *R23E10* ∩ *VGlut-GAL4*-driven expression of *VGlut*^RNAi^ or EKO reduces daily sleep (left, *P*<0.0001, Dunn’s test after Kruskal-Wallis ANOVA) without altering waking locomotor activity (right, *P*>0.9999 relative to ≥1 parental control, Dunn‘s test after Kruskal-Wallis ANOVA). The left panel is reproduced from Fig. 3i. **d**, *R23E10* ∩ *VGlut-GAL4*-driven expression of *VGlut*^RNAi^ reduces the time courses and percentages of sleep regained after deprivation by vortex (SD) (left panels, genotype effects: *P*≤0.0314, time × genotype interactions: *P*<0.0001, two-way repeated-measures ANOVA; right panel, *P*≤0.0429, Dunn’s test after Kruskal-Wallis ANOVA). **e**, Sleep profiles of flies expressing CsChrimson under the control of *R23E10 ∩ R84C10*, with or without retinal (*n*=35 and 36 flies, respectively), before, during, and after optogenetic replay of SWA (20 light pulses s^-1^ in 500-ms bursts) (normalized sleep: 1.31±0.07, retinal effect: *P*=0.0032, time × retinal interaction: *P*<0.0001, two-way repeated-measures ANOVA). **f**, **g**, *R23E10 ∩ R84C10*-driven expression of *VGlut*^RNAi^ reduces daily sleep (**f**, left, *P*≤0.0209, Holm-Šídák test after ANOVA) and the time courses and percentages of sleep regained after deprivation (SD) (**g**, left, genotype effect: *P*=0.0039, time × genotype interaction: *P*<0.0001, two-way repeated-measures ANOVA; right, *P*≤0.0251, Dunn’s test after Kruskal-Wallis ANOVA) without altering waking locomotor activity (**f**, right, *P*≥0.1481 relative to ≥1 parental control, Dunn‘s test after Kruskal-Wallis ANOVA). Data are means ± s.e.m.; *n*, number of flies; asterisks, significant differences (*P*<0.05) in planned pairwise comparisons. For statistical details see Supplementary Table 3.

**Figure S 6.**
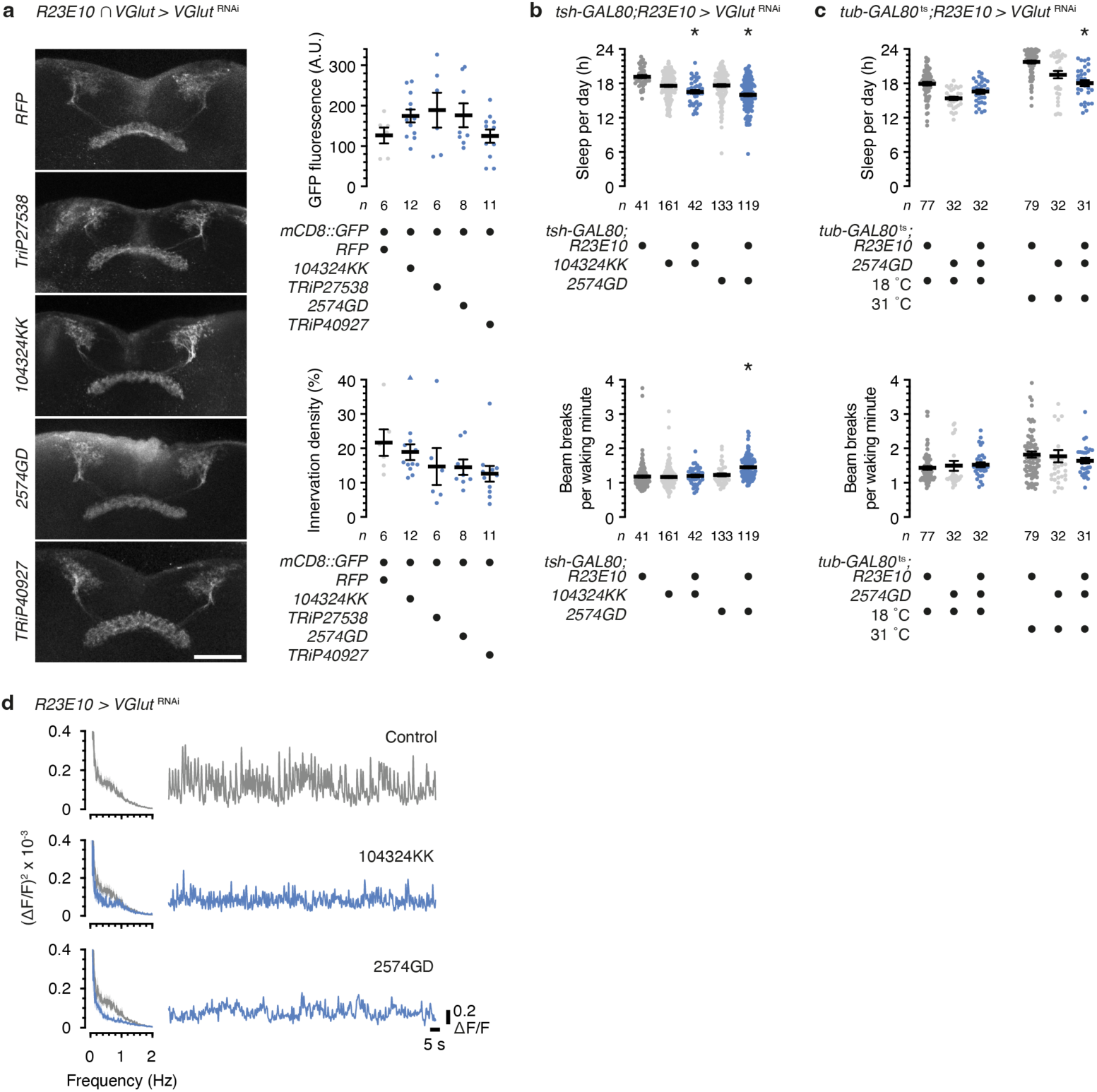
Chronic and acute interference with glutamatergic transmission from dFBNs reduces sleep and SWA without altering neuronal morphology. **a**, Maximum-intensity projections of dFBNs in flies expressing RFP or *VGlut*^RNAi^ under the control of *R23E10* ∩ *VGlut-GAL4* (left). Scale bar, 50 μm. GFP fluorescence (top right) and innervation density (bottom right) of dFBN projections is unaffected by the expression of *VGlut*^RNAi^ (*P*≥0.0608, Dunn’s test after Kruskal-Wallis ANOVA). One data point exceeding the y-axis limit is plotted as a triangle at the top of the innervation density graph; mean and s.e.m. are based on the actual values. **b**, *R23E10*-driven expression of *VGlut*^RNAi^ reduces daily sleep in the presence of teashirt-GAL80 (top, *P*≤0.0174, Dunn’s test after Kruskal-Wallis ANOVA). Expression of *VGlut*^RNAi^ ^2574GD^ (*P*≤0.0001), but not of *VGlut*^RNAi^ ^104324KK^, increases waking locomotor activity (bottom, *P*≥0.5732, Dunn’s test after Kruskal-Wallis ANOVA). **c**, Temperature-inducible, adult-specific expression of *VGlut*^RNAi^ under the control of *R23E10-GAL4* reduces daily sleep (top, temperature × genotype interaction: *P*=0.0035, two-way ANOVA) without altering waking locomotor activity (bottom, temperature × genotype interaction: *P*=0.3797, two-way ANOVA). **d**, Power spectra (left) and example dendritic GCaMP traces (right) of dFBNs expressing GCaMP without or with *R23E10-GAL4*-driven *VGlut*^RNAi^. Data are means ± s.e.m.; *n*, number of flies; asterisks, significant differences (*P*<0.05) in planned pairwise comparisons. For statistical details see Supplementary Table 3.

**Figure S 7.**
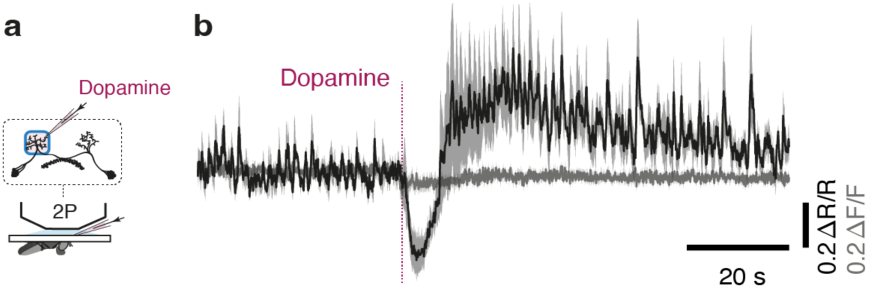
Dopamine inhibits dFBNs. **a**, Imaging of dFBN dendrites during pressure ejection of 10 mM dopamine. **b**, Average GCaMP trace (black) normalized to mCD4::tdTomato fluorescence (ΔR/R) and aligned to a 640-ms dopamine pulse. Average ΔF/F of mCD4::tdTomato is shown for comparison (grey). *n*=8 ROIs in 4 flies. Data are means ± s.e.m. For imaging details see Supplementary Table 1.

**Figure S 8.**
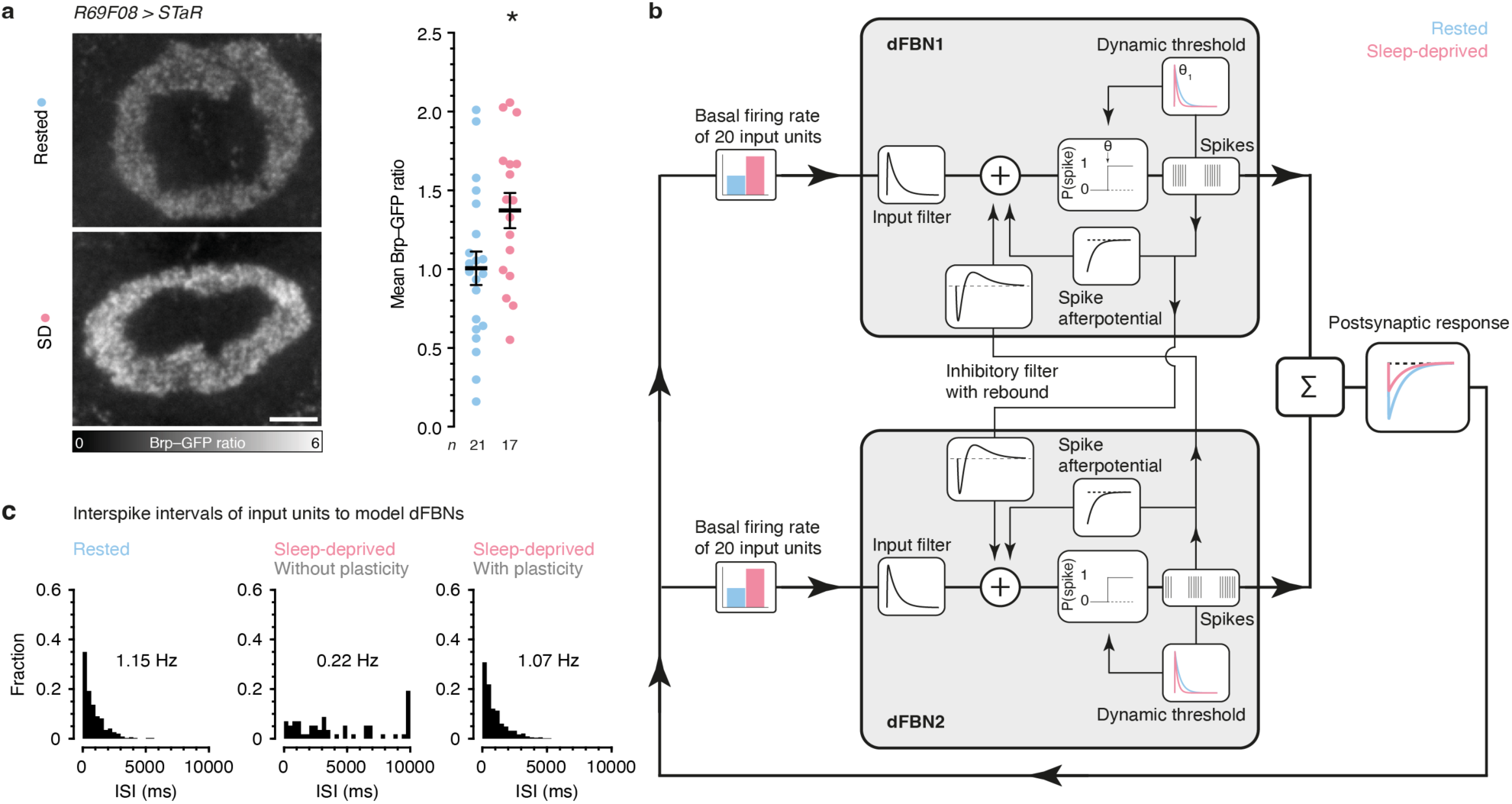
Efferent dFBN synapses depress after sleep deprivation. **a**, Summed-intensity projections of V5-tagged endogenous BRP in R5 neuron axons coexpressing mCD8::GFP. Emission ratios are intensity-coded according to the key at the bottom and increase after sleep deprivation (SD) (*P*=0.0236, *t* test). Scale bar, 10 μm. **b**, Model architecture. dFBN1 and dFBN2 are each driven by 20 presynaptic units whose spike trains are inhomogeneous Poisson processes; the rates of these Poisson processes are subject to feedback inhibition by the summed spikes of both dFBNs. The membrane potential of each dFBN is a linear sum of resting potential, excitatory postsynaptic potentials due to Poisson units (input filter), inhibitory postsynaptic potentials and postinhibitory rebound due to the contralateral dFBN (inhibitory filter with rebound), and spike afterpotential. The membrane potential passes through a dynamic threshold to produce spikes. Color indicates sleep history-dependent variations in the basal firing rates of Poisson units, the dynamic threshold of dFBNs, and the postsynaptic response to transmission from dFBN synapses. **c**, Interspike interval (ISI) distributions of Poisson units at baseline and after sleep deprivation, in the absence and presence of plastic feedback connections from dFBNs. Data are means ± s.e.m.; asterisk, significant difference (*P*<0.05) in a planned pairwise comparison. For statistical details see Supplementary Table 3.

**Figure S 9.**
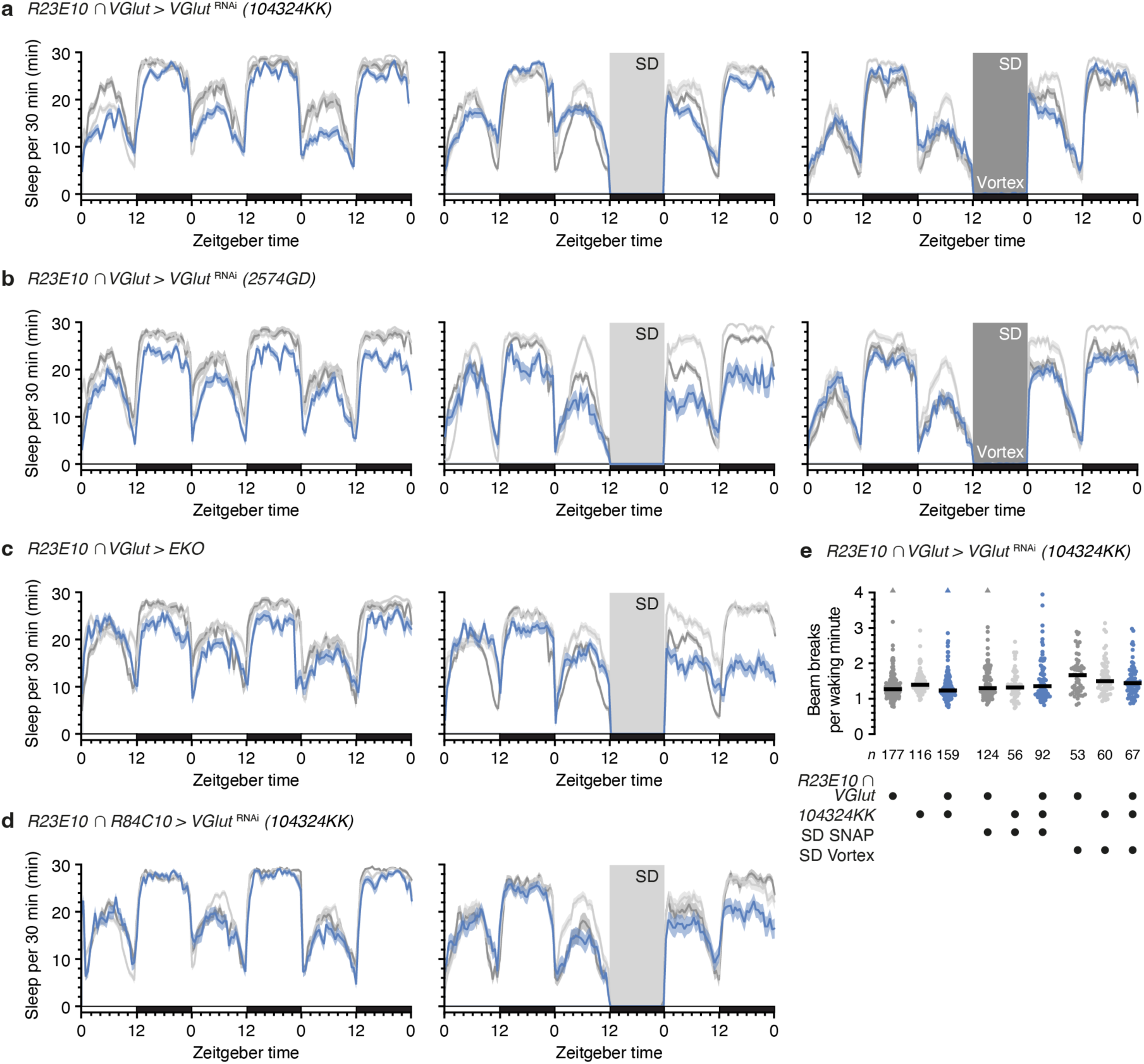
Sleep profiles during unperturbed sleep, two forms of sleep deprivation, and recovery. **a**–**d**, Sleep profiles of experimental flies (blue) expressing *VGlut*^RNAi^ (**a**, **b**, **d**) or EKO (**c**) under the control of *R23E10* ∩ *VGlut-GAL4* (**a, b, c**) or *R23E10* ∩ *R84C10-GAL4* (**d**) and parental controls (grey). Sample sizes and symbol colors as in Fig. 3 and Fig. S 5, which report summary data. Left column, unperturbed sleep; center column, 12-h sleep deprivation (SD) on day 2 via the sleep-nullifying apparatus (SNAP); right column, 12-h sleep deprivation (SD) on day 2 by vortex. Two-way repeated-measures ANOVA detects significant genotype effects on sleep (*P*≤0.0056) and significant time × genotype interactions (*P*<0.0001) during all quantification periods (left column, days 2 and 3; center and right columns, day 3). **e**, Waking locomotor activity in flies expressing *VGlut*^RNAi^ under the control of *R23E10* ∩ *VGlut-GAL4* and parental controls, before and after sleep deprivation (SD) by SNAP or vortex. Three data points exceeding the y-axis limit are plotted as a triangles at the top of the graph; mean and s.e.m. are based on the actual values. Mildly elevated waking activity after sleep deprivation rules out injury (effect of sleep history: *P<*0.0001, effect of SD method: *P*=0.2287, genotype effect: *P*=0.5877, three-way ANOVA). Data are means ± s.e.m.; *n*, number of flies. For statistical details see Supplementary Table 3.

**Figure S 10.**
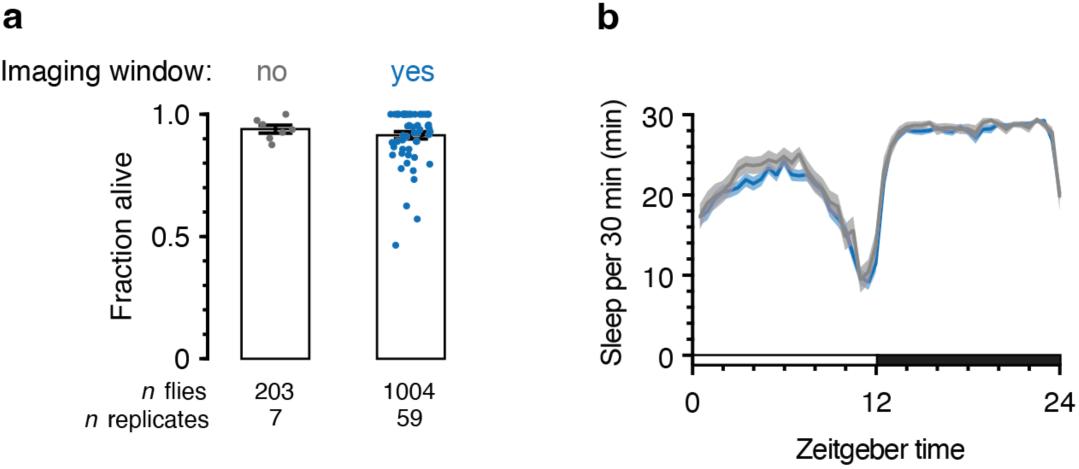
Viability and sleep profiles of flies implanted with a chronic optical imaging window. **a**, Implantation of a chronic optical imaging window does not reduce the survival of operated flies at 24–48 h after surgery (blue) relative to controls (grey) (*P*=0.9874, Mann-Whitney test). **b**, Sleep profiles of flies implanted with a chronic imaging window (blue, *n*=74) vs. controls (grey, *n*=37). Two-way repeated-measures ANOVA fails to detect significant effects on sleep (effect of window implantation: *P*=0.2355, time × window implantation interaction: *P*=0.9890).

## Legends to Supplementary Videos

**Supplementary Video 1.** Simultaneous two-photon imaging of GCaMP signals in dFBN dendrites of both hemispheres. The video, which is thresholded for display, was acquired at a frame rate of 14.56 Hz in a continuous imaging session lasting ∼22 minutes.

**Supplementary Video 2.** Two-photon imaging of GCaMP signals in dFBN somata. The video, which is thresholded for display, was acquired at a frame rate of 14.56 Hz in an imaging session lasting 96 s.

**Supplementary Video 3.** Confocal image stack of the CNS of a fly expressing 6xEGFP under the control of *R23E10-GAL4* in the presence of the transcriptional repressor VGlut-GAL80. Green, 6xEGFP; blue, BRP. Scale bars, 100 μm.

**Supplementary Video 4.** Confocal image stack of the CNS of a fly expressing 6xEGFP under the control of hemidrivers *R23E10-DBD* and *VGlut-p65AD* (*R23E10 ∩ VGlut-GAL4*). Green, 6xEGFP; blue, BRP. Scale bars, 100 μm.

---

**Supplementary Table 1.** Two-photon imaging parameters.

| Experiment and figures | N pixels (rows x columns) | Axial coordinates of focal planes ($\mu\text{m}$)† | Acquisition rate (Hz)† | Sealed imaging window* | Zeitgeber time | Additional details |
| --- | --- | --- | --- | --- | --- | --- |
| Dendritic GCaMP imaging (Fig.1a, 2, 4, Fig. S1f,g,h, 2) | 256 x 256 | 0, 10, 30, 40 | 14.56 | Yes | 0–4<sup>(+)</sup>; specified in Fig. 4. | ROI selection blind to sleep history and zeitgeber time. Data displayed in Fig.1a, 2, 4, Fig. S,1f-h and Fig. S1 are from the same dataset. (20 $\mu\text{m}$ instead of 40 in vortex experiments) |
| Cell body GCaMP imaging (Fig. 1b,c) | 256 x 256 | Four planes at variable distances of 0-30 $\mu\text{m}$ | 14.56 | No | 2–13 | No between-group comparisons and thus no blinding during ROI selection. In some cases data were obtained from the left and right hemispheres of the same brain in separate imaging sessions. These were treated as independent experiments. dFBNs with a slow rhythm: 0–2 per hemisphere (median: 1). |
| Simultaneous cell body and dendritic GCaMP imaging (Fig. S1a,b,c,d,e) | 256 x 256 | Four planes at variable distances of 0-40 $\mu\text{m}$ | 14.56 | No | 2–17 | No between-group comparisons and thus no blinding during ROI selection. In some cases data were obtained from the left and right hemispheres of the same brain in separate imaging sessions. These were treated as independent experiments. |
| Simultaneous patch-clamp and GCaMP imaging (Fig.1d,e,f) | 256 x 256 | 0 | 58.25 | No | 3–12 | No between-group comparisons and thus no blinding during ROI selection. |
| Dendritic GCaMP imaging plus VGlut RNAi (Fig.3k, Fig. S 6d) | 256 x 256 | 0, 10, 20, 30 | 14.56 | Yes | 1–9 | ROI selection blind to genotype. |
| iGluSnFR, iAChSnFR, and GRAB<sub>ACh</sub> imaging during optogenetic stimulation of dFBNs (Fig.3d) | 100 x 256 | 0, 10 | 68.32 | No | 4–11 | No blinding during ROI selection. Stimulus-aligned traces are means of 20 repetitions per fly, applied every 6 s. |
| GCaMP imaging during optogenetic stimulation of a dFBN subset (Fig.3e) | 256 x 256 | Two planes at distances of 7-13 $\mu\text{m}$ | 29.13 | No | 3–11 | ROI selection blind to dietary retinal and genotype. Stimulus-aligned traces are means of several simultaneously recorded cell bodies per fly, averaged over 10 repetitions (light applied every 20 s; 1-2, 1-3, and 1-5 cell bodies in flies expressing CsChrimson::tdTomato with retinal, CsChrimson::tdTomato without retinal, and no CsChrimson::tdTomato, respectively. In some cases data were obtained from the left and right hemispheres of the same brain. These were recorded in separate imaging sessions and averaged post hoc. Average number of identified CsChrimson::tdTomato-positive cells per fly: 2.2 ± 0.92 (mean ± standard deviation). |
| GRAB<sub>DA2m</sub> imaging during heat stimulation (Fig.5b) | 256 x 256 | 0, 10, 20, 30 | 14.56 | Yes | 1–10 | No between-group comparisons and thus no blinding during ROI selection. |
| Dendritic GCaMP imaging during heat stimulation (Fig. 5c,d,e) | 256 x 256 | 0, 10, 20, 30 | 14.56 | Yes | 3–11 | No between-group comparisons and thus no blinding during ROI selection. |
| GRAB<sub>DA2m</sub> imaging during optogenetic stimulation of dFBNs (Fig.5g) | 256 x 256 | 0, 10, 20, 30 | 14.56 | Yes | 1–12 | ROI selection blind to genotype. |
| iGluSnFR imaging during optogenetic stimulation of dFBNs after sleep deprivation (Fig.6b) | 100 x 256 | 0, 10 | 68.32 | Yes | 0–5 | ROI selection blind to sleep history. Stimulus-aligned traces are means from two simultaneously recorded axonal ROIs (different focal planes) and 5 repetitions (for each light intensity) per fly. The order in which stimuli with different intensities were applied was varied between flies to avoid adaptation. Flies were placed on retinal food after optical window implantation and kept on retinal food during sleep measurements. |
| GCaMP or iAChSnFR imaging during focal application of dopamine (Fig. S7) or acetylcholine (Fig. 3d inset), respectively | 256 x 256 | Four planes at distances of 0-50 $\mu\text{m}$ | 14.56 | No | 4–10 | No between-group comparisons and thus no blinding during ROI selection. |

† In ascending order from ventral to dorsal.
¶ Volumetric rate when more than one focal plane was imaged.
\* If no sealed imaging window was used, dissections were done on the recording day and brains were superfused with extracellular solution (see Methods).

**Supplementary Table 2.**
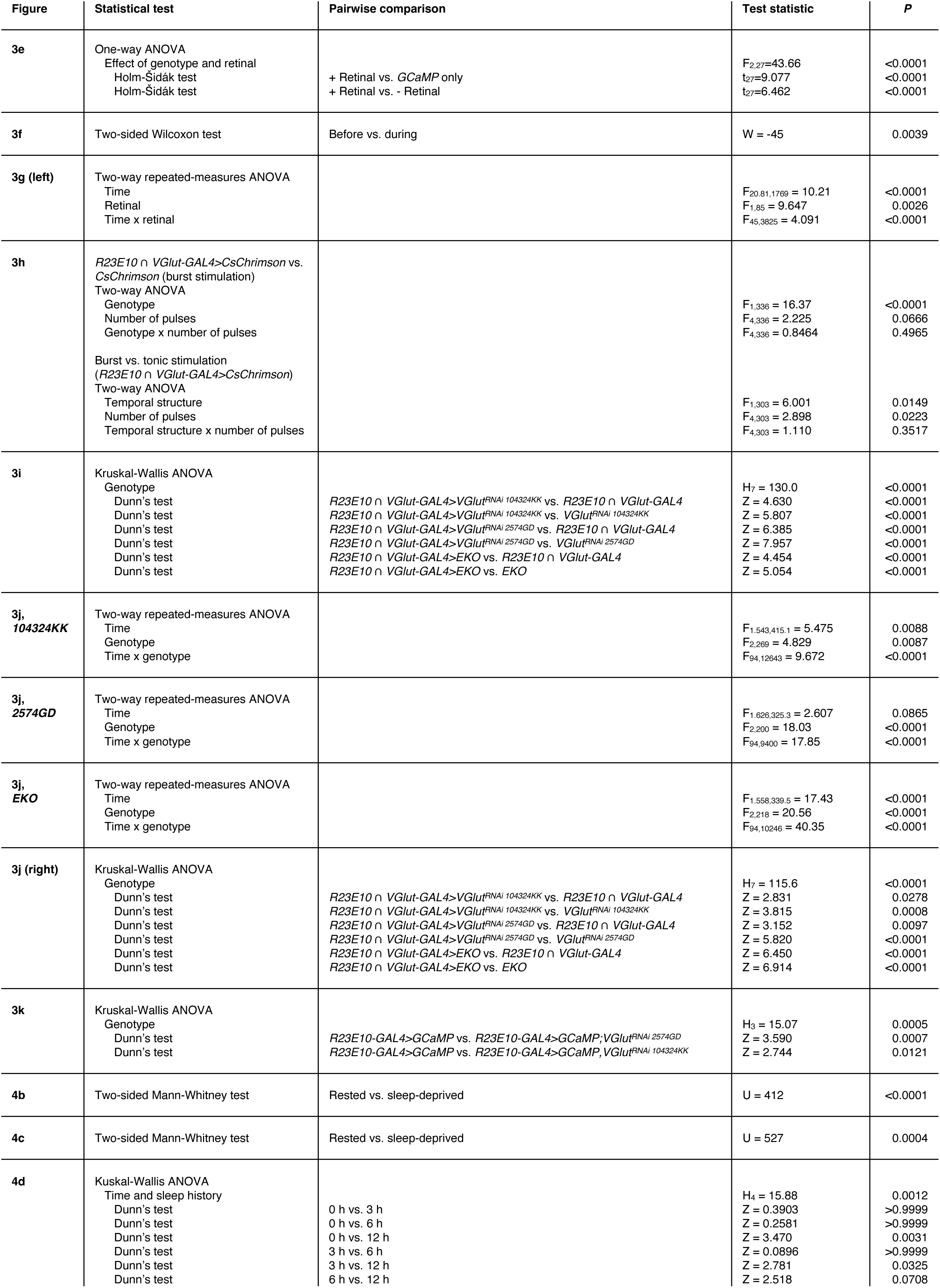
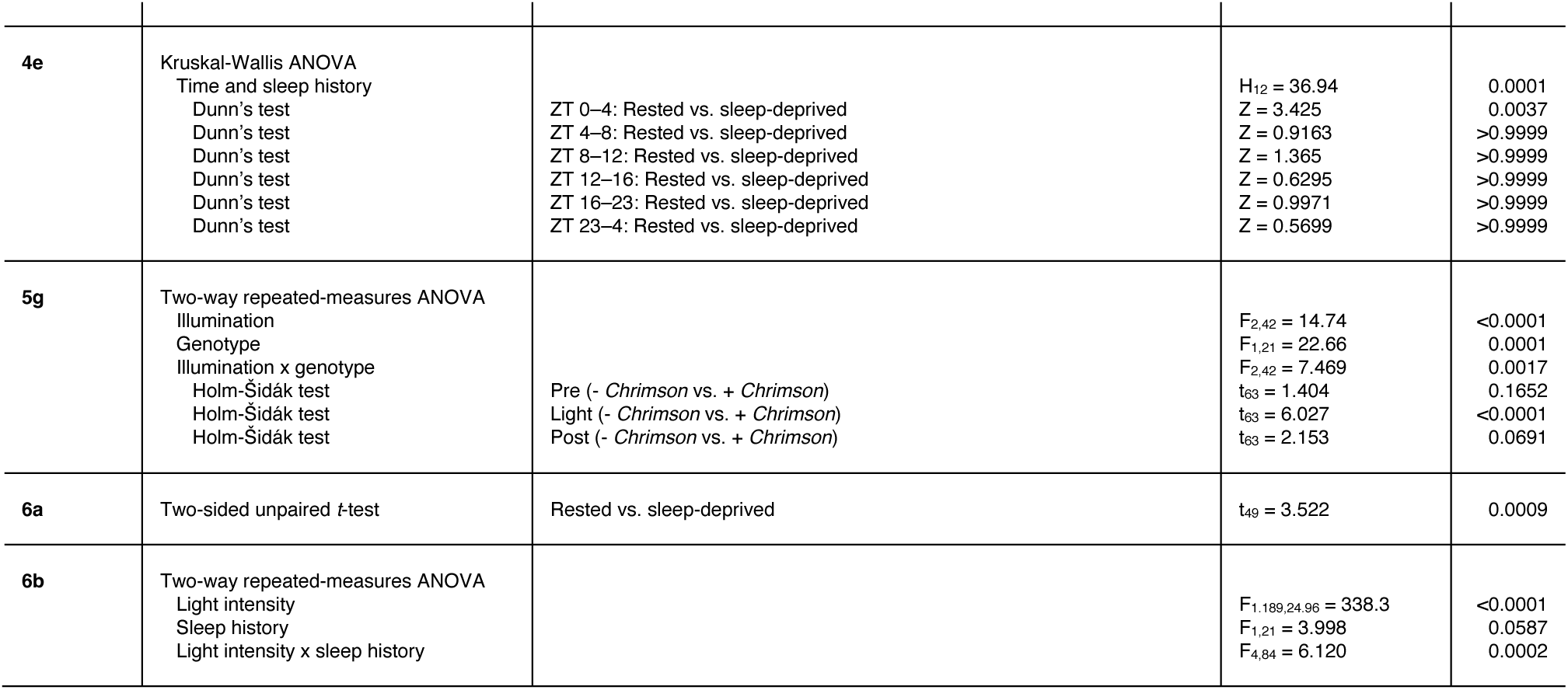
Statistical analyses of Figures 3–6.

**Supplementary Table 3.**
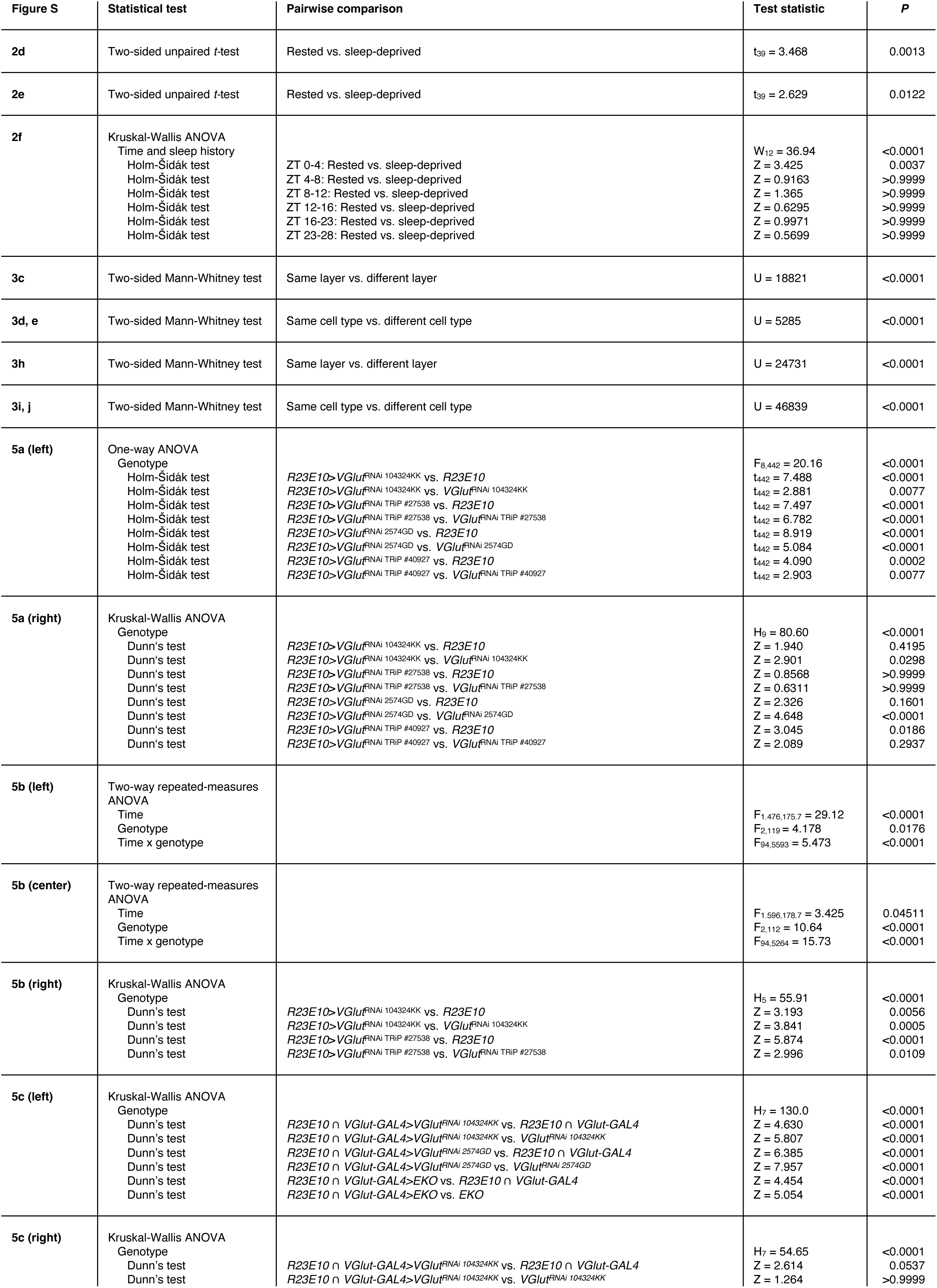
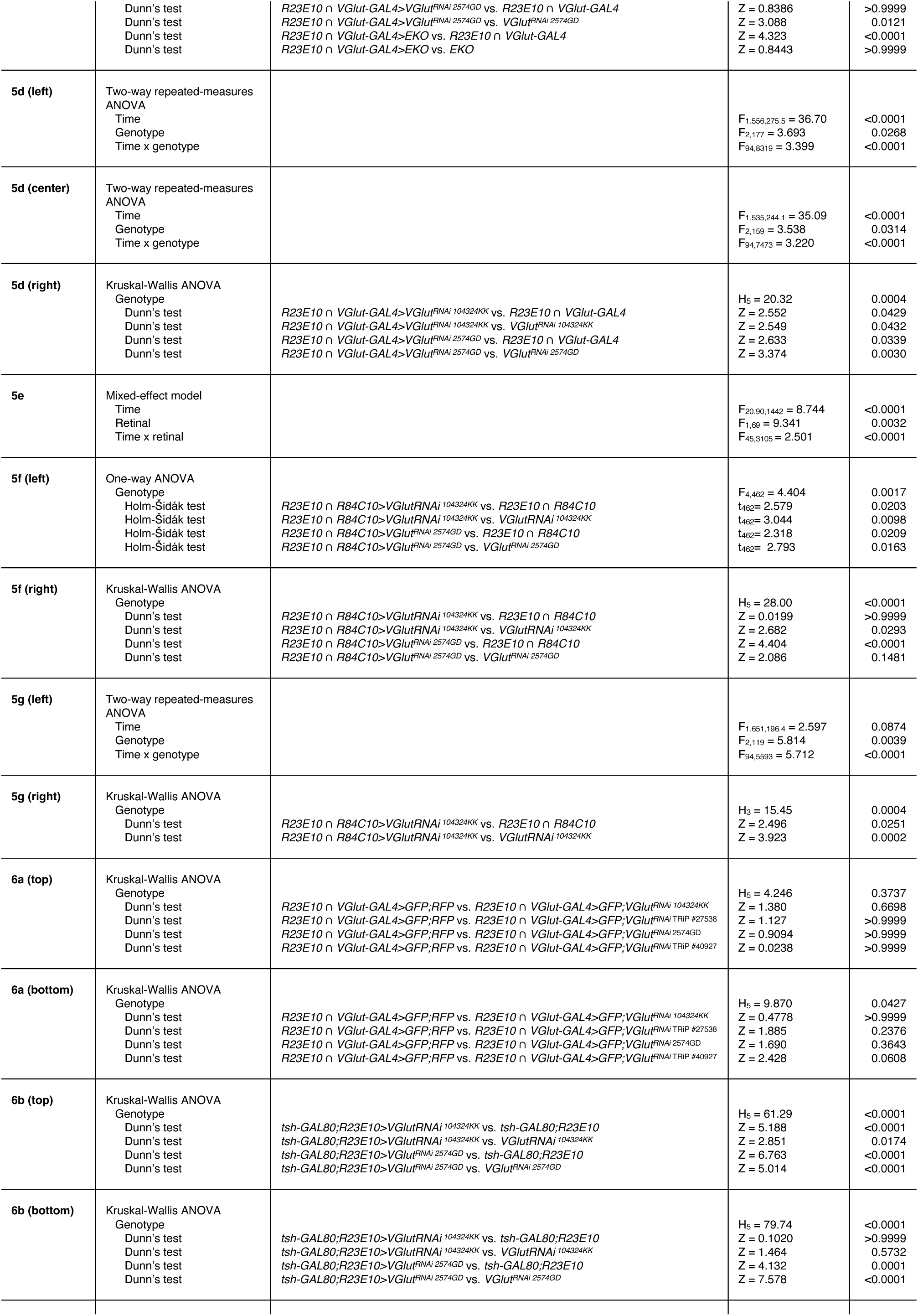
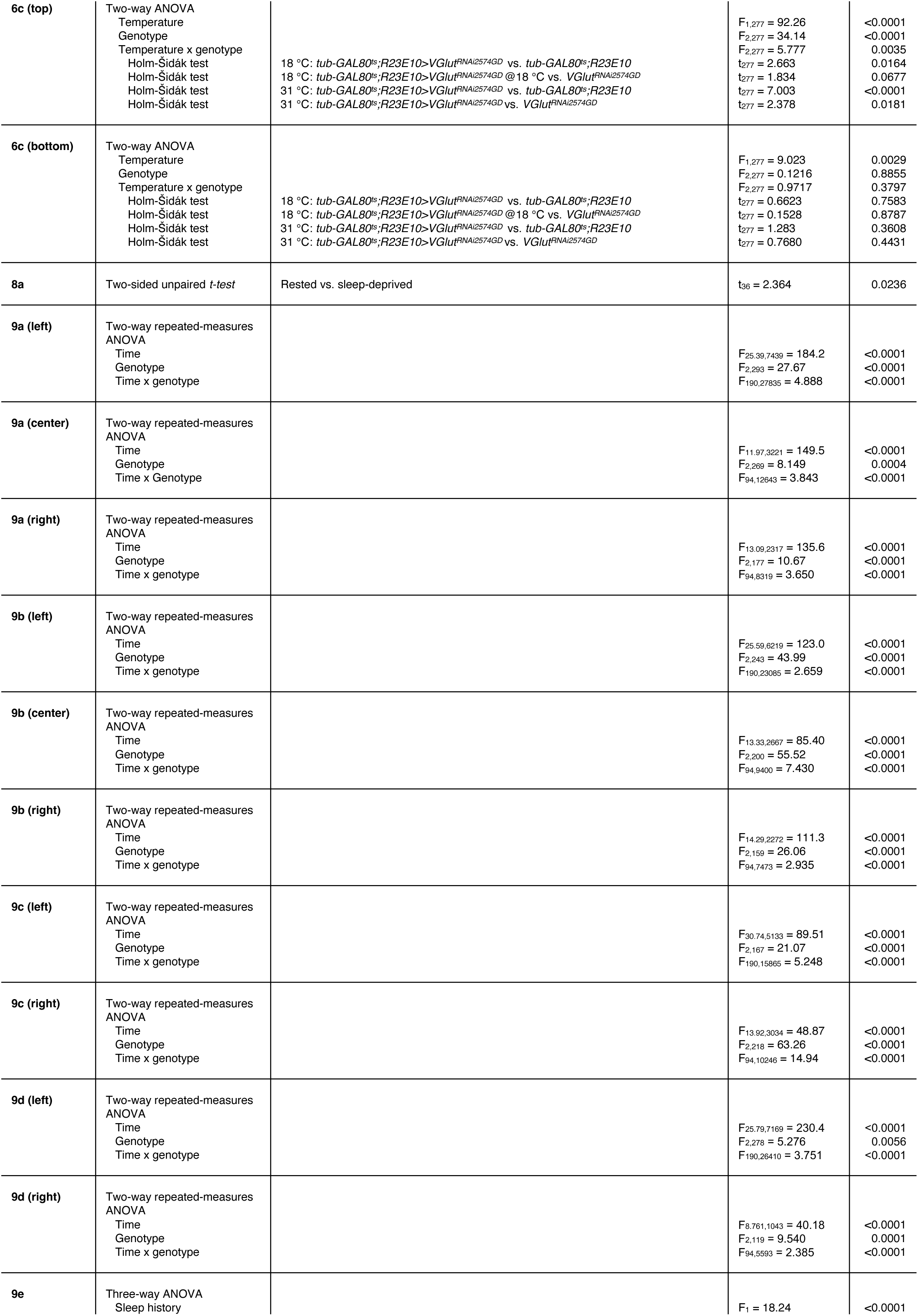
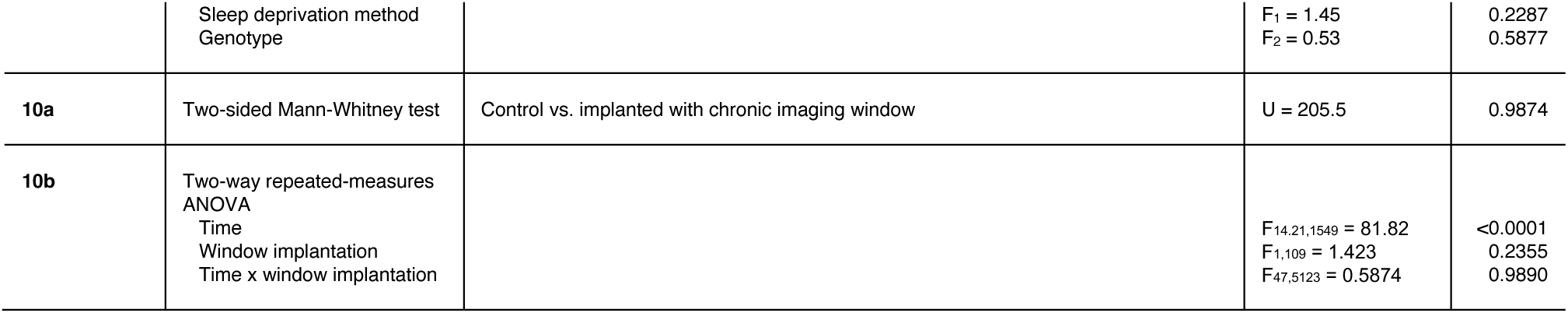
Statistical analyses of Figures S 2–10.

